# Profiling of yeast *Saccharomyces cerevisiae* mitochondrial AMPylome reveals a regulation of ATP synthase coupling trough subunit δ

**DOI:** 10.1101/2024.12.14.628500

**Authors:** Chiranjit Panja, Suchismita Masanta, Aneta Wiesyk, Katarzyna Niedzwiecka, Emilia Baranowska, Dominik Cysewski, Marta Sipko, Anna Anielska-Mazur, Agata Malinowska, Roza Kucharczyk

**Affiliations:** Institute of Biochemistry and Biophysics, Polish Academy of Sciences, Warsaw, Poland

## Abstract

The adenylation (AMPylation) of proteins as a posttranslational modification is used by bacteria during infection of host cells. These new virulence factors - AMPylases mainly belonging to the FIC domain containing proteins and constitute a potential drug target. Human FIC protein (HYPE) controls the activity of BiP chaperone under endoplasmic reticulum stress. No FIC family proteins have yet been identified in yeast *Saccharomyces cerevisiae*. The second family of AMPylases are SelO proteins which control the redox homeostasis in mitochondria and chloroplasts. We describe here the first global screening of AMPylated proteins in yeast *S. cerevisiae* mitochondrial proteome from wild type and SelO (Fmp40) lacking cells. Through quantitative mass-spectrometry-based proteomics, we identified a total of 169 AMPylated proteins in mitochondria while AMPylated peptides of 115 proteins were identified in *fmp40Δ* mitochondria, indicating on the presence of another, besides Fmp40, not yet identified AMPylase in yeast. We confirmed AMPylation of Atp1, Atp2, Atp3 and Atp16 subunits of mitochondrial ATP synthase by western blotting. Interestingly, we found AMPylation and phosphorylation of many residues, what indicates on the complex regulation of the ATP synthase activity. We confirmed the importance of one of such residues in Atp16, showing that its post-translational modification serves to regulate ATP synthase and OXPHOS coupling in both fermentative and respiratory growth conditions. This regulation serves to maintain the proper potential of the inner mitochondrial membrane, particularly under conditions of fermentative growth. This dataset represents the first library of AMPylated mitochondrial yeast proteins reported to date and supplements the AMPylome of human chronic lymphocytic leukemia cell line from human HYPE containing and HYPE lacking cells. The data represents a foundation for substrate specific investigations that can ultimately decipher the biological role of the AMPylation in the mitochondria.

## INTRODUCTION

Mitochondria fulfil multiple functions in the cell, the most important of which is ATP synthesis by oxidative phosphorylation (OXPHOS), a process catalyzed by the respiratory enzymes organized in supercomplexes in the mitochondrial inner membrane [1, 2]. During this process, electrons from carbohydrates and fatty acids are transferred to oxygen, which results in a proton gradient across the mitochondrial inner membrane that is used by the ATP synthase to phosphorylate ADP. This activity is very important to make a proper polarization of the inner mitochondrial membrane (ΔΨ). Beyond ATP synthesis, mitochondria are critical for numerous metabolic pathways, like synthesis of amino acids, lipids, heme, and iron-sulfur clusters and are the source of internal signals, that direct the cell to programmed cell death [3–5]. In addition, mitochondria have their DNA (mtDNA), encoding several subunits of the OXPHOS, therefore, their biogenesis, which depends on the energy demand of the cell, requires the coordination of gene expression from nuclear and mitochondrial genomes and the import of the mitochondrial proteins from the cytoplasm to the organelle [6, 7]. The functioning of the OXPHOS is accompanied by the continuous production of reactive oxygen species (ROS), the levels of which perform important signaling functions in the cell and are strictly controlled [8, 9]. All above processes must be tightly regulated to permit mitochondria to respond to changes in energy demand, changes in cellular metabolism, or environmental conditions [10, 11]. The mitochondrial and cellular functions are controlled at the level of gene expression (in a coordinated fashion between two genomes) - mRNA synthesis and turnover, the coordination of protein synthesis and degradation, and activity regulation by posttranslational modifications (PTM). The best studied PTM is phosphorylation of hydroxyl-containing amino acids (serine, threonine and tyrosine) by the addition of a phosphoryl (PO_3_^−^) group by protein kinases [12]. Studies by many groups focused on the regulation of the mitochondrial proteome by phosphorylation have shown that 67 percent of the mitochondrial proteome is phosphorylated with an average of 6.76 modified residues per protein [13–18]. For some of these modifications, their biological significance has been described in the literature, including Atp2 and Atp20 subunits of ATP synthase, and the Qcr2 subunit of the complex III [13, 19–21].

The same amino acid residues, which are phosphorylated, are subject to another modification - the covalent attachment of adenosine monophosphate (AMP), called AMPylation or adenylation. This modification was first discovered in 1967 as a mechanism controlling the activity of bacterial glutamine synthetase [22–24]. AMPylation was for the first time found in eukaryotes in the context of bacterial infection when Yarbrough *et al.* reported AMPylation of host small GTPases by bacterial virulence factor VopS from *Vibrio parahemeolyticus*. AMPylation precludes interactions of GTPases with downstream binding partners and causes actin cytoskeleton collapse leading to host cell death [25]. The bacterial AMP transferases have been classified as either filamentation induced by cyclic AMP (FIC) or adenylyl transferase (AT) domain containing enzymes [23, 26]. In humans, there is one FIC domain containing AMPylase, FicD (also known as HYPE or hFic, encoded by *FICD* gene), localized in the endoplasmic reticulum, modulating the unfolded protein response by AMPylation of the HSP70 family chaperone BiP [27–29]. In 2018 another novel class of AMPylases have been discovered: proteins of Selenoprotein O (SelO) family, classified to the pseudo-kinases [30, 31]. Mammalian and *S. cerevisiae* SelO (Fmp40) are localized in the mitochondria, are regulated by its redox state and are active when are reduced [31–33]. SelÒs biological role is to control the redox homeostasis in this organelle by regulating the global protein glutathionylation [31]. Yeast Fmp40 level is induced upon respiration and exposition to hydrogen peroxide, mitochondrial redoxins Prx1, Trx3 and Grx2 are AMPylated by Fmp40 *in vitro* and physical interaction of Fmp40 with Trx3 and Prx1 was demonstrated *in vivo* [34]. Fmp40 regulates the redox cycles of Prx1 and Trx3 and one of the mechanisms is AMPylation of T66 residue of Trx3, which is crucial for the proper Trx3 protein level and maturation during its import to the mitochondrial matrix [34]. Fmp40 is the only AMPylase identified in yeast to date and it was postulated that this organism does not have the FIC domain containing protein [29]. A few SelO substrates in *A. thaliana* and *E. coli* have been identified by immunoprecipitation and mass spectrometry approaches, and the full range of SelO targets in other model organisms is yet to be determined [31, 35].

Few approaches aiming to identify the AMPylated proteins were performed to date. The first work described the chemo-proteomic screening and substrate validation for HYPE-mediated AMPylation in mammalian cell lysate and identified 25 AMPylated proteins [36]. The second work used antibodies against adenosine-phosphate followed by anti-AMP-Tyr to purify AMPylated proteins from the human chronic lymphocytic leukemia *FICD* wild-type and *FICD* knock-out cell lines extracts [37]. A total of 213 proteins were identified from wild type cell extracts, while only 23 of these were pulled down from HYPE lacking cells, consistent with the presence of another AMPylase, besides HYPE in mammalian cells.

This study presents the first comprehensive analysis of the mitochondrial AMPylome within native mitochondria isolated from *S. cerevisiae*. Employing both global and affinity purification approaches targeting mitochondrial ATP synthase, Trx3-myc redoxin, and Fmp40-GFP AMPylase, we identified 169 AMPylated proteins and 318 distinct AMPylation sites in a high confidence mitochondrial proteome. Two-dimensional BN-PAGE/SDS-PAGE electrophoresis with AMP-specific antibodies further confirmed AMPylation of ATP synthase subunits. In parallel, we conducted an analysis of mitochondrial protein phosphorylation, revealing many novel phosphorylation sites not previously documented in yeast mitochondrial phosphoproteome. Newly identified phosphoproteins include OXPHOS complex subunits (Qcr10, Cox5a, Cox9, Cox13, Cox2, Coq1), the mitochondrial import machinery component Tim23, and mitochondrial thioredoxin Trx3, suggesting a greater extent of mitochondrial protein phosphorylation than previously recognized. Importantly, we demonstrate a critical role for Atp16-S29 that can be posttranslationally modified by both AMPylation and phosphorylation, in maintaining mitochondrial inner membrane potential, particularly during growth on fermentable carbon sources.

## Materials and methods

### Yeast strains, media, and growth conditions

The MR6 strain (*MATa ade2-1 his3-1,15 leu2-3,112 trp1-1 ura3-1 arg8∷HIS3*) used in the study is a derivative of W303-1B strain background (Table 1) [38]. Strain expressing Atp6-HA-6His was a gift from prof. Alexander Tzagoloff (Columbia University, NY, USA, [39]). The strain expressing the *TRX3-MYC* gene was a kind gift from Prof. Chris Grant (University of Manchester, UK, [40]). Strains were grown in rich YPGA medium (1% Bacto yeast extract, 1% Bacto peptone, 2% glucose, 40 mg/L adenine), YPGlyA medium (1 % Bacto yeast extract, 1 % Bacto peptone, 2 % glycerol, 40 mg/L adenine) or SC selective medium: (6.7 g/L YNB w/o amino acids, 2% glucose or glycerol, 40 mg/L adenine, appropriate amino-acids drop-out powder (Sunrise)) at 28°C with shaking at 180 rpm.

**Table 1.**
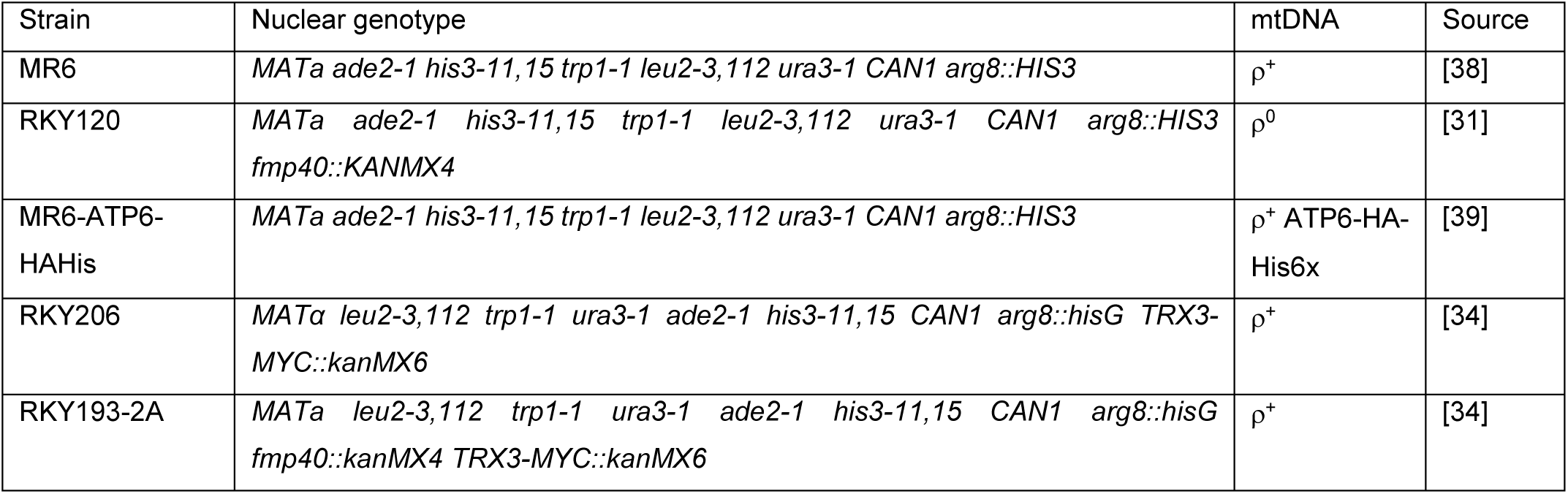
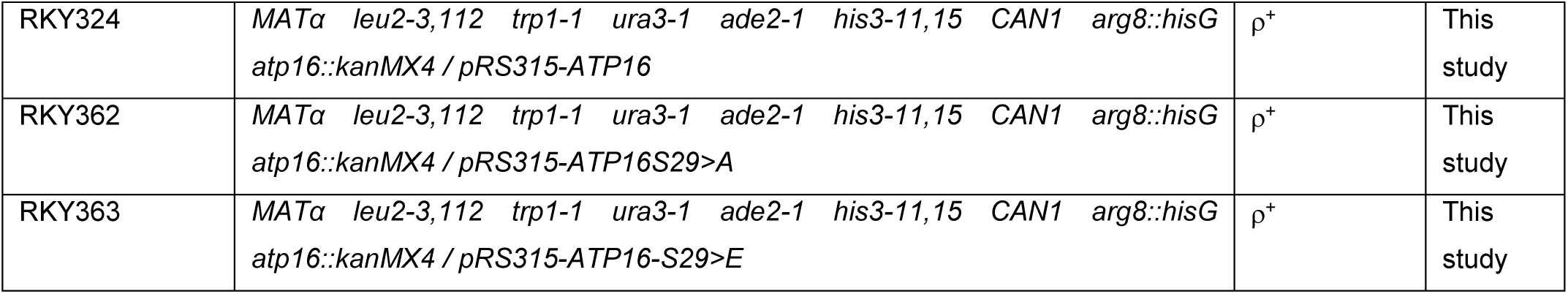
Genotypes and sources of yeast strains.

The Atp16, Atp16-S29>A and Atp16-S29>E were expressed from the centromeric pFL38 plasmid (the pFL38-ATP16 was provided by dr Michaela Carraro, [41]. The site-directed mutagenesis was performed according to the protocol of Q5 Site-Directed Mutagenesis Kit (*NEBiolabs*) using this plasmid as a template with the pairs of primers:

ATP16-S29>A 5‘GTTCATATGCAGAAGCTGCTGCCGCA**GCA**TCAGGTTTGAAGTTACAATTTGC3’ or ATP16-S29>E 5‘GTTCATATGCAGAAGCTGCTGCCGCA**GAA**TCAGGTTTGAAGTTACAATTTGC3’ and ATP16-S29short 5‘ GCTTAGCGACGAAATTCAATG3’. The correct sequence of *ATP16* gene variants in resulting plasmids, pRK98 and pRK99, respectively, were verified by sequencing. The plasmids were introduced into the wild type MR6 strain. Such a strains were transformed with a *atp16::KANMX4* deletion cassette, amplified with ATP16-Up and ATP16-Low primers (5‘TGCCCAGCCAATCAAAGCGC3’, 5‘CGATGATCTGCAACTAGGGC3’, respectively) using the total DNA from *atp16::KANMX4* knock-out strain obtained from Open Biosystems. Transformants were selected on the YPGA + geneticin (200 µg/ml) plates. The correct integration of the cassette into the genomic *ATP16* locus in RKY324, RKY362 and RKY363 (Table 1), not the plasmid-born copies, was verified by PCR with primers ATP16-low and Kan-dln (5‘GATTTTGATGACGAGCGTAATG), because the primer ATP16-low sequence is absent in the plasmid pFL38-ATP16.

### Isolation of pure mitochondria for mass spectrometry (MS) analysis

Mitochondria were isolated from cells grown in YPGlyA at 28°C by the enzymatic method according to the protocol described previously with minor modifications [42, 43]. Briefly, cells grown to OD600 of 4 were centrifuged, washed with cold water, and incubated for 10 min in the SH buffer (0.1 M Tris pH 9.3, 0.5 M 2- β-mercaptoethanol) at 32°C. After washing twice with the KCl buffer (10 mM Tris pH 7.0, 0.5 M KCl) cells were digested by Zymolyase T20 (ref. 07663-91, Nacalai Tresque, Japan) at 8 mg/1 g of dry weight calculated from the equation (number of liters x OD of the culture x 0.28) in the 10 ml / 1 g of dry weight Digestion buffer (1.35 M sorbitol, 1 mM EGTA, 10 mM citric acid, 30 mM disodium phosphate) for 30 min at 32°C. Spheroplasts were washed twice with the Washing buffer (10 mM Tris-Maleate pH 6.8, 0.75 M sorbitol, 0.4 M mannitol, 0.1 % BSA), suspended in 40 ml of Homogenization buffer (10 mM Tris-Maleate pH 6.8, 0.6 M mannitol, 2 mM EGTA) and homogenized 3 x 5 sec. in the Mixer blender in the 37 ml inox cup (WARING, #W67902, #W87278S). After centrifugation of the cell debris at 1076 x g for 8 min, the homogenate was centrifuged at 11952 x g for 10 min at 4°C (Sorval RC 5BPlus, SS-34 rotor). Mitochondria were washed once with the Homogenization buffer and suspended in 9 ml of 15% Percoll in the Homogenization buffer. The mitochondria were placed on the 23%-40% Percoll-Homogenization buffer gradient and purified by ultracentrifugation in Sorvall WX Ultra Series Centrifuge (Thermo Fisher Scientific) at conditions: 31176 x g (28000 rpm), 14 min, 4°C, acceleration 2, deceleration 1, T-8100 rotor). The pure mitochondrial fraction was picked from 23%-40% phase interface and washed twice in Homogenization buffer with the centrifugation in Eppendorf 5804R centrifuge at 15557 x g, 20 minutes, 4°C, F34-6-38 rotor. Mitochondria were suspended in 250 µl of sonication buffer (100 mM Tris-HCl pH 7,4; 6 M guanidine hydrochloride, inhibitors of proteases cOmplete Mini (ref. 11836153001, Roche, Indianapolis, IN), inhibitors of phosphatases Phosphatase Inhibitor Cocktail I Liquid (ref. J63907, Thermo Fisher Scientific) and sonicated in Bioruptor XL Diagenode 30 times 30 sec at “high” with 30 sec intervals on ice. Proteins were precipitated using cold acetone (1 ml pre-cooled acetone was added to the sample and incubated for 1 hour at-20°C, then harvested using OHAUS FC5515R centrifuge at 9500 x g, 10 min at 4°C, the protein pellets were dried at 37°C for 30 min, and analyzed by mass spectrometry. The experiment was repeated ten times.

### Purification of ATP synthase complexes and Trx3-Myc from mitochondria

To pull-down the ATP synthase monomer the strain expressing Atp6 subunit C-terminally tagged by HA-6xHis from the mitochondrial genome was used [44]. The whole ATP synthase complexes were purified from 5 mg of mitochondria by Ni-NTA agarose beads as described in reference [45]. The 2 µg of Ni-NTA bead eluates were loaded on the 15 % SDS-PAGE gel. Then the gel was stained with Coomassie blue or silver staining according to manufacturer’s protocol (Pierce sliver stain kit, Thermo Fisher Scientific) to visualize the proteins before their MS analysis. The experiment was done in triplicate.

The ATP synthase dimers and the monomers were extracted from 1 mg of the mitochondria divided into 5 tubes in extraction buffer (30 mM HEPES pH 6.8, 150 mM potassium acetate, 12% glycerol, 2 mM 6- aminocaproicacid, 1 mM EGTA, 1.5 % digitonin (Sigma), protease inhibitors (ref. 11836153001, Roche, Indianapolis, IN), by incubation on ice for 26 min. The samples were centrifuged using OHAUS FC5515R centrifuge at 21950 x g, 4 °C, 30 min, supplemented with 4.5 μL of loading dye (5 % Serva Blue G-250, 750 mM 6-aminocaproicacid), run on NativePAGE^TM^ 3–12 % Bis-Tris Gels (Thermo Fisher Scientific). Dimers and monomers were cut off from the gel and analyzed by MS. The experiment was done in triplicate.

For Trx3-Myc and Fmp40-Myc purification, the RKY206 (Table 1) or wild type and RKY120 transformed with pUG35 based plasmid pSP2 (encoding Fmp40-Myc C-terminal fusion) were used. 1 mg of mitochondria isolated from cells grown on rich or minimal selective lacking uracil, glycerol containing media, was centrifuged using OHAUS FC5515R centrifuge at 9500 x g for 10 minutes and suspended in 400 µL of the solubilization buffer (20 mM Tris/HCl pH 7.4, 0.1 mM EDTA, 50 mM NaCl, 10% [vol/vol] glycerol, 1% [wt/vol] digitonin, 2 mM PMSF, 1 x EDTA free proteinase inhibitor (Roche, Indianapolis, IN)) and incubated for 30 minutes on ice. After centrifugation as above, the supernatant was collected and incubated with 200 µL of Pierce Anti-c-Myc Magnetic beads (Thermo Fisher Scientific). The bound proteins were analyzed by MS. The above experiments were done in duplicate.

### 2^nd^ dimension electrophoresis of ATP synthase complexes

For second-dimension electrophoresis, the dimers and the monomers were cut off from the first BN-PAGE gel, and incubated for 10 min in the SDS-PAGE electrophoresis buffer supplemented with β-mercaptoethanol (50 µL of per 10 mL of buffer), placed into the lanes of the Novex™ Tris-Glycine Mini Protein 16% gel (Thermo Fisher Scientific). Proteins were transferred to the cellulose membrane with the iBlot2 Gel Transfer Device (Thermo Fisher Scientific) and followed by Western blotting with polyclonal antibodies against Atp2, Atp3, Atp4, Atp5, Atp16 (a kind gift from dr Marie-France Giraud, Bordeaux), and anti-Tyr-AMP (ref. ABS184, Merck) and anti-Tre-AMP (ref. 09-890, Merck).

### Mass spectrometry analysis

The tryptic digested peptides were analyzed by LC/MS system at the IBB PAS Mass Spectrometry Facility unit using Evosep One (Evosep Biosystems, Odense, Denmark) coupled to a Orbitrap Exploris 480 mass spectrometer (Thermo Fisher Scientific, Bremen, Germany) via Flex nanoESI ion source (2) or by a nanoACQUITY UPLC system (Waters, Milford, MA) coupled to either Orbitrap Velos (Thermo Fisher Scientific, Bremen, Germany) or Q Exactive (Thermo Fisher Scientific, Bremen, Germany) mass spectrometer. Data were gathered in a data-dependent manner in a positive mode. Raw files were later analyzed with the Proteome Discoverer (version 2.4.1.15, Thermo) and Sequest search engine. First, data was searched against the SGD database (version 20210423) supplemented with the database of popular contaminants (CRAP), with the following parameters – enzyme: semiTrypsin, missed cleavages: 2, precursor mass: 10 ppm, fragment mass: 0.02 Da, variable modifications: Oxidation (M), Phosphoadenosine (HKSTY), Phosphorylation (STY). Fixed modification depended on sample preparation method and was defined either as Carbamidomethyl (C), Methylthio (C) or none. Search results were later validated with Percolator, with target FDR for PSM, peptide and protein level equal 0.01 [46]. The results included only high confidence modification sites and peptides (high confidence in at least 1 sample). Data were also pre-processed with Mascot Distiller (version 2.8, Matrixscience) and searched with Mascot search engine (version 2.8, Matrixscience) against the SGD database, with offline mass re-calibration (with MScan software, version 3.0, https://proteom.ibb.waw.pl/mscan/) based on results of preliminary Mascot searches, typical resulting parent mass window of 5 ppm and fragment mass window of 0.01 Da. The rest of the search parameters were similar to the ones used in Proteome Discoverer searches. Results were validated using target/decoy strategy, with threshold q value of 0.01.

### Functional enrichment of proteins

To investigate the functional roles of the identified proteins, pathway and process enrichment analysis were performed using Metascape with enrichment analysis conducted against a background within the yeast genome. Significantly enriched terms were identified based on a p-value and the significant terms were grouped into clusters based on their shared gene memberships. To further elucidate the interrelationships between these terms, a subset of 20 clusters were chosen by prioritizing terms with the most significant p-values within each of the 20 clusters, with a maximum of 15 terms per cluster and a total limit of 250 terms was chosen for visualization. The resulting network was visualized using Cytoscape, with nodes representing enriched terms and color-coded based on their respective cluster IDs. To further investigate the enriched pathways of the identified proteins, protein interaction networks were constructed using STRING database with medium confidence level [47] and visualized using the STRING 106 plugin in Cytoscape. The identified proteins were classified manually or by Gene Ontology terms.

### Measurement of oxygen consumption, ATP synthesis and membrane potential

Mitochondria were isolated from cells grown in SC-glycerol-ura medium at 28°C according to the protocol described above, but without their purification by percol gradient centrifugation. For all assays, they were diluted to 75 µg/ml in respiration buffer (10 mM Tris-maleate pH 6.8, 0.65 M mannitol, 0.35 mM EGTA, and 5 mM Tris-phosphate). Oxygen consumption rates were measured using a Clarke electrode adding consecutively 4 mM NADH (state 4 respiration), 150 µM ADP (state 3) or 4 µM carbonyl cyanide m-chlorophenylhydrazone (CCCP) (uncoupled respiration), as described previously [48]. Complex IV activity was measured by first adding to mitochondria and CCCP to uncouple respiration. Then, simultaneously 100 µM ascorbate and 3 µM N,N,N′,N′-Tetramethyl-p-phenylenediamine (TMPD) were added and oxygen consumption rates measured. The rates of ATP synthesis were determined under the same experimental conditions with 750 µM ADP. Every 15 seconds, 100 µl aliquots were withdrawn from the oxygraph cuvette and added to 50 µl of the 3.5 % (w/v) perchloric acid and 12.5 mM EDTA solution already prepared in the tubes to stop the reaction. The samples were then neutralized to pH 6.5 by the addition of KOH and 0.3 M MOPS. The synthetized ATP was quantified using a luciferin/luciferase assay (Kinase-Glo Max Luminescence Kinase Assay, Promega) in a Beckman Coulter Paradigm plate reader. The participation of F1F_O_-ATP synthase in ATP production was assessed by measuring the sensitivity of ATP synthesis to oligomycin (3 μg/ml). Variations in transmembrane potential (ΔѰ) were evaluated by monitoring the fluorescence quenching of Rhodamine 123 (0.5 μg/mL; λ_exc_ of 485 nm and λ_em_ of 533 nm) from mitochondrial samples (0.150 mg/mL) in the respiration buffer under constant stirring at 28°C using a Cary Eclipse Fluorescence Spectrophotometer (Agilent Technologies, Santa Clara, CA, USA) as described previously [49].

### Mitochondrial membrane potential measurement *in vivo*

Yeast cells were grown to logarithmic phase overnight in SC-glucose-ura 28°C. 2 OD of yeast cells were pelleted down, washed with medium, and incubated in the same growth media containing 100 nM of MitoTracker Red CMXRos (Life Technologies) for 30 min at room temperature. Half of the control cells culture was passed to another tube and CCCP was added to 25 µM final concentration. After 10 minutes of shaking the MitoTracker Red CMXRos was added without washing off CCCP. After staining cells were spun down and resuspended in the same media, and only one picture was taken immediately by fluorescence microscopy, as the fluorescence signal dropped very fast with time. This fact resulted in multiple technical repetitions until the appropriate number of photographed cells for statistics was obtained [50]. All experiments were performed with four biological replicates. Images were collected using Nikon C1 confocal scanning head mounted on an Eclipse TE2000-E inverted microscope equipped with CFI Plan Apochromat VC 100x oil objective (NA 1.4). MitoTracker CMXRos fluorescence was excited with green light emitted by a He-Ne Laser 543 nm laser (Melles Griot,1.0 mW) set at 3% then collected with a 610 long pass emission filter and displayed in false 16 colours (LUTs). In all experiments the acquisition settings like the laser power, the gain of the PMT, the pixel dwell, the pinhole size was the same. Reference DIC images were collected concomitantly by the detector. Stack of confocal sections (each consist of 10 optical sections with the 0.5 µm intervals) were collected in Nikon EZ-C1 software. The collected z-stacks were processed by 3D Deconvolution module in NIS-AR 6.02.03 (Nikon) and digitally processed using FIJI software (NIH, Bethesda, MD, USA). All presented images are the maximum intensity z-projections and were compiled in in FigureJ plugin (NIH, Bethesda, MD, USA; https://journals.plos.org/plosone/article?id=10.1371/journal.pone.0240280). To compare levels of the mitochondrial membrane potential in tested strains the fluorescence intensity was quantified using FIJI (Fiji Is Just ImageJ 2.14.0; Java 1.8.0-322 [64-bit]). Z-max projections obtained from the deconvoluted stacks were segmented using Otsu’s automatic thresholding method. The accuracy of the mask was visually checked to ensure that the selected area corresponded to the mitochondria and additional manual selection was introduced. Mean fluorescence intensity was measured in the individual region of interest (ROI) and specifies accurately the net of mitochondria per cell. For each strain and CCCP-treatment the 60 cells were quantified. Kruskal-Wallis statistical tests were performed with GraphPad Prism (10.4.0; San Diego, CA, USA) and the statistical details are included in the figure legends.

## Data and Statistical analyses

Unless otherwise stated in the figure legends or method section, each experiment was repeated at least three times. Data are presented as a representative experiment or as the average ± s.d. To assess significant differences with the control and test samples, the unpaired student’s t-test was used. The structure of Atp16 was analyzed from AlphaFold Protein Structure Database (AF-Q12165-F1-v4) and the structural visualization was carried out using PyMOL software. Images were processed using Adobe Photoshop (Adobe Inc.) and figures were assembled using Microsoft Office PowerPoint. The proteomic data are available via ProteomeXchange with identifier PXD058585.

## Results

### Defining a high confidence yeast mitochondrial proteome

The mitochondria of high purity from wild type and *fmp40Δ* strains grown in the non-fermentative carbon source glycerol, were obtained by differential centrifugation and subsequent percoll gradient centrifugation [51]. To preserve the AMPylation sites the mitochondrial proteins were precipitated with cold acetone, dried and analyzed by nano-LC-MS/MS. The ATP synthase dimers and monomers were either separated in the native gels or subjected to affinity chromatography (using the strain expressing *ATP6-HA- 6xHis* from the mitochondrial genome) and subsequently subunits of purified enzyme were separated by SDS-PAGE (Figs. 1A and S1). The bands containing the ATP synthase dimers, monomers or subunits and co-purified associated proteins were cut off from the gels, subjected to in-gel digestion, and analyzed by nano-LC-MS/MS. Additionally, we purified proteins associating with Fmp40-Myc (expressed from the plasmid) and similar approach was used to purify Trx3-Myc mitochondrial redoxin (expressed from the genome), as we have found that this redoxin interacts with Fmp40 *in vivo* [34]. To define a high confidence mitochondrial proteome, we first screened and compared our identified proteins with previously reported mitochondrial proteins from six large scale studies that aimed to identify the total yeast mitochondrial proteome [52–57] and the list of curated yeast mitochondrial proteins present in SGD database. In addition, we analyzed the identified proteins using three different predictive analysis determining mitochondrial protein localization and presence of mitochondrial import sequences [58–60]. This gave us a total of 1322 proteins. We further considered proteins to be authentic mitochondrial proteins if they were absent in the literature or previous works but were identified both in Percoll purified mitochondrial proteins and in any other analysis of proteins from ATP synthase dimer and monomer from native gels or Trx3-Myc pull down. However, only proteins identified only in Percoll purified mitochondria or in any other pull-down analysis but not in Percoll purified mitochondria were ignored. This gave an additional 398 proteins which made the total number of proteins identified in our analysis to be present in the yeast mitochondria to 1720 (Table S1). The proteins were classified using Metascape and the most enriched clusters are used for visualization using Cytoscape software (Fig. 1B).

**Figure 1.**
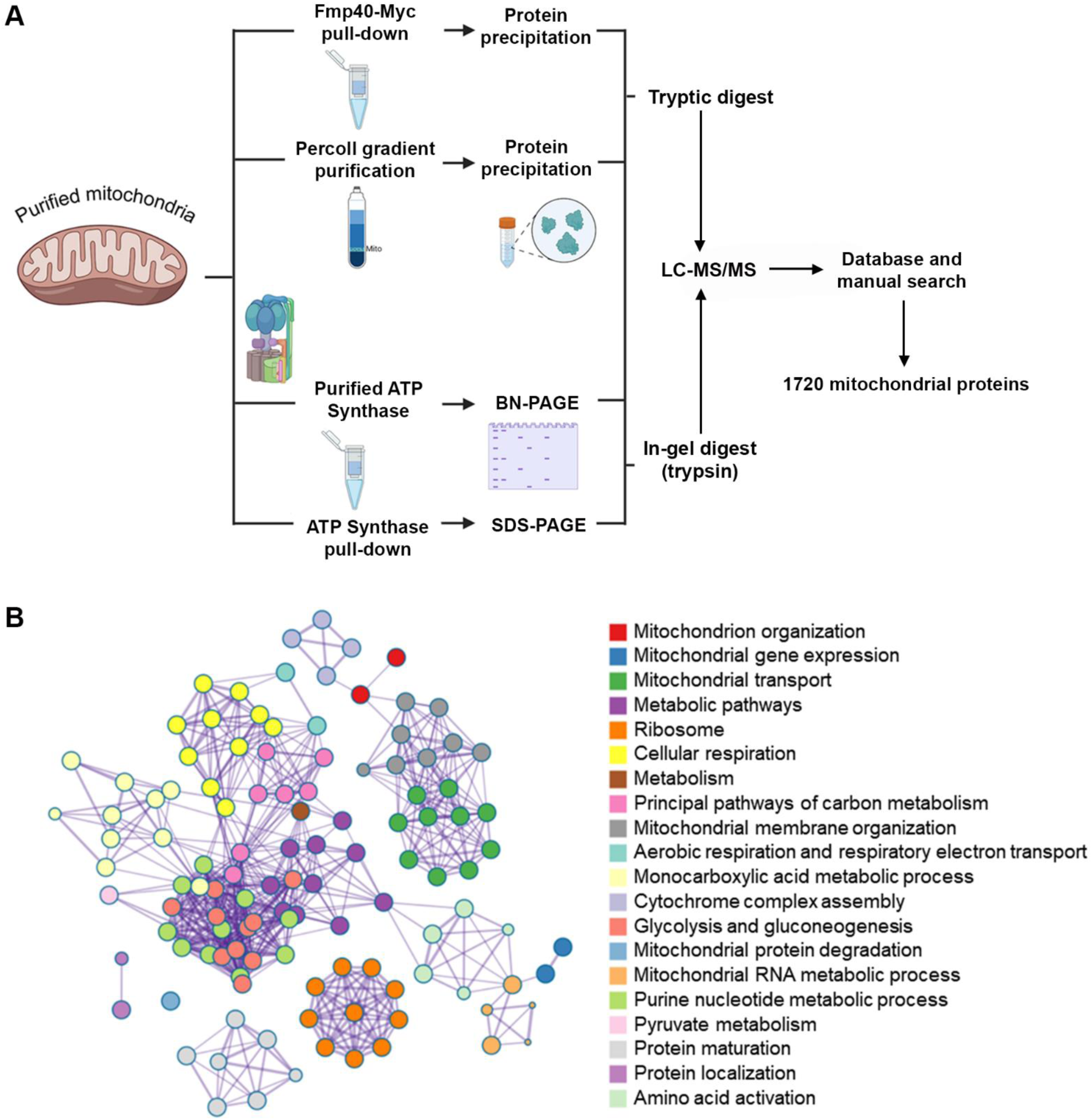
Scheme of experimental approaches for identification of mitochondrial proteins. **A** Starting with yeast mitochondria, several parallel approaches are outlined in the figure: purification of mitochondria by percoll gradient, pull down of Fmp40 or Trx3 and associated proteins, pull-down of ATP synthase and SDS-PAGE separation of its subunits, and extraction of ATP synthase complexes for BN-PAGE separation and subsequent LC-MS/MS analysis. The technical details are described under “Material and methods”. **B** Functional classification of identified mitochondrial proteins using Metascape. The top 20 most significantly enriched clusters are depicted in the cluster with terms exhibiting high similarity score being connected by edges in the network.

### Identification of AMPylation Sites in Yeast Mitochondrial Proteins

Using strict criteria for the interpretation and validation of AMPylated peptide spectra as outlined under “Materials and methods,” we identified 318 different AMPylation sites in 296 peptides originating from 169 proteins from the identified mitochondrial proteome in this study. In 103 proteins a single AMPylation site was found. In 32 proteins – two, in 17 proteins – three, and in 7 proteins – four AMPylation were identified. We found five or more AMPylation sites in eleven proteins: five in Atp16, six in Ilv5, Cit1, Atp1 and Tef2, seven in Mia40, eight in Por1 and, nine in Ssc1 and Hsp60 and ten in Atp2. We found 83 AMP-serine (S) residues, 70 AMP-threonine (T) residues, 22 AMP-tyrosine (Y) residues, 22 AMP-histidine (H) residues, and 101 AMP- lysine (K) residues modifications with high confidence. In 21 cases the exact position of the AMPylated residue could not be precisely assigned by the software due to several potential modification sites present in a peptide and insufficient fragmentation data. 107 peptides were present only in samples from the wild-type strain, absent in *fmp40Δ* cells – among them 36 modifications of serine (43 % of all serine modifications), 27 modifications of threonine (39% of all threonine modifications), and 7 modifications of tyrosine residues (32% of all tyrosine modifications). These modifications, potentially dependent on Fmp40 AMPylase, are among others in Atp1, Atp2, Atp7, Atp16, Coq5, Eno2, Hsp60, Idh1, Kgd1, Kgd2, Mdl2, Mia40, Mir1, Mss4, Nde1, Ndi1, Sam2, Ssa2, Ssb2, Ssc1, Tdh3, Tef2, Trx3, Vps1, Yml6, and Vps7 proteins. However, ∼60 % of serine/threonine/tyrosine residues modifications were present in samples from *fmp40Δ* cells, indicating the presence of other AMPylases in yeast mitochondria specific to these residues, independently on the AMPylases specific to histidine or lysine residues. The identified AMPylation sites are summarized in Table 1 and Supplemental Table S2.

### Functional Classification of Identified AMPylated Proteins

We grouped the identified AMPylated proteins according to their determined or proposed functions (Fig. 2) based on gene ontology curation. The two largest groups were formed by proteins involved in mitochondrial energy and carbon metabolism and metabolic pathways. We identified 38 proteins which were known to bind ATP to be AMPylated. Among them, the most significant group of proteins that were AMPylated belongs to heat shock proteins and proteins involved in protein folding pathway (Ecm10, Hsp60, Hsp78, Ssa1, Ssa2, Ssb2, Ssc1, Ssq1, and Mdj1). The other major group of proteins found to be AMPylated in the mitochondrial proteome were those belonging to the oxidoreductase family including ten members of the Flavohemoproteins family, NAD(P) binding proteins and other dehydrogenases, indicating the redox proteins are particularly susceptible to AMPylation in the mitochondria. Remarkably, fifteen subunits of the OXPHOS including three subunits of Complex III (Qcr2, Qcr6 and Qcr8), one subunit of Complex IV (Cox6) and seven subunits of the ATP synthase complex (Atp1, Atp2, Atp3, Atp5, Atp7, Atp15, Atp16) were found to be AMPylated. Thus, AMPylation is observed in a broad spectrum of yeast mitochondrial proteins.

**Figure 2.**
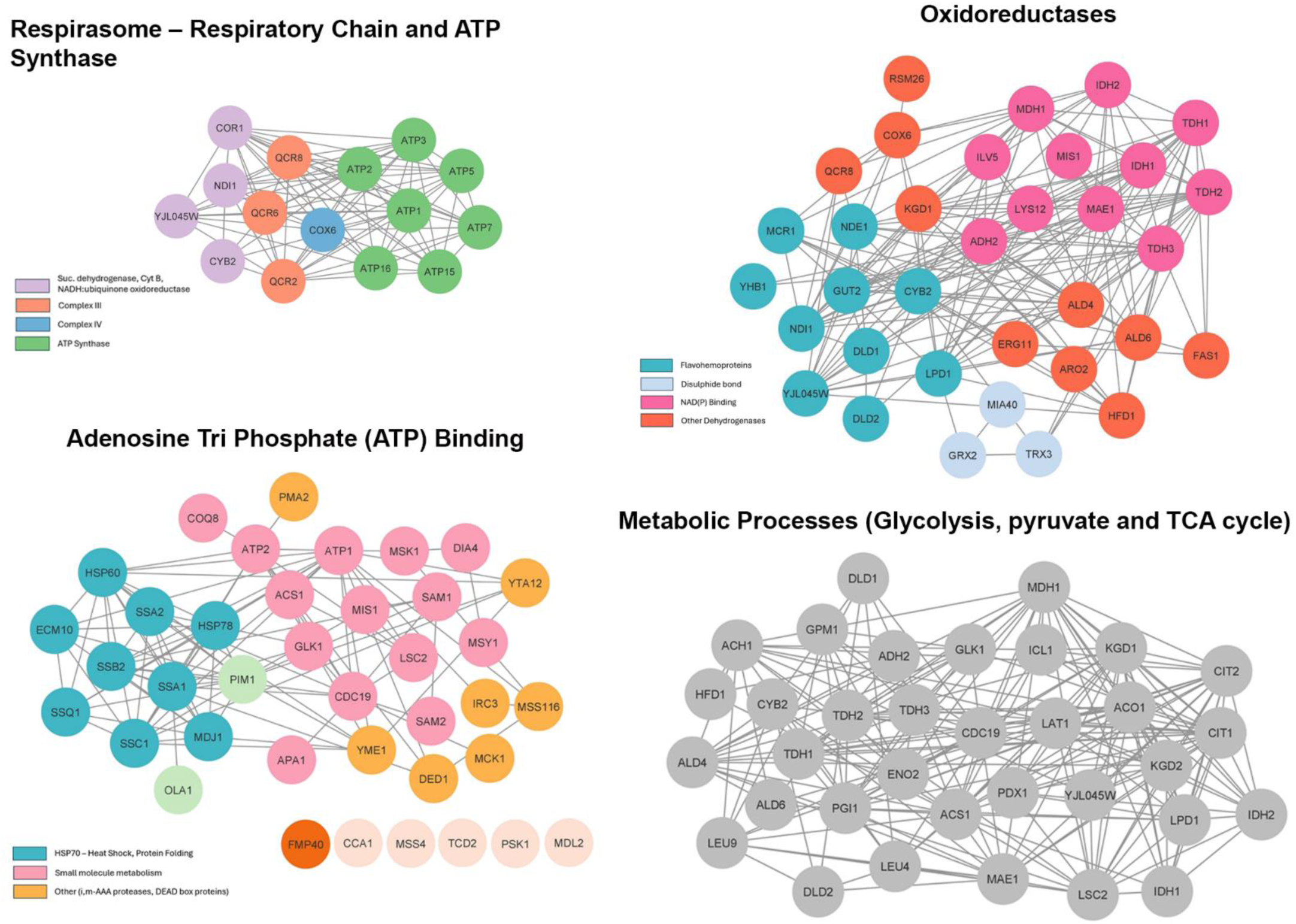
Functional enrichment of AMPylated proteins in mitochondria. 169 identified mitochondrial AMPylated proteins from the total yeast mitochondrial proteome. The most enriched Gene ontology clusters were created using STRING database and visualized using Cytoscape.

### New Phosphorylation Sites of Mitochondrial Proteins

The raw data from all the experiments were also analyzed to identify the phospho-peptides. We found 1540 phosphorylation sites among 516 proteins (Table S3). The list of all phosphorylation sites present in our experiments was compared to the phospho-sites reported previously and present in the yeast protein phosphorylation database YeastMine (https://yeastmine.yeastgenome.org/yeastmine, updated on 15 July 2024, [61]). The list of 1077 phosphorylation sites not detected in previous experimental approaches is included in Table S3 and it is 2.4 % of the already reported phospho-sites in the yeast proteome. Of the 1720 proteins defined to be mitochondrial in this study, we found 1216 phosphorylation sites in 332 proteins, among those sites 823 are reported for the first time. Focusing on the OXPHOS enzymes, redox enzymes or subunits of mitochondrial protein import systems, several of them have not been previously found in the phospho-proteome. Among them are Qcr10, Cox5a, Cox9, Cox13, Cox2, Coq1, Trx3, or Tim23, indicating that yeast phospho-proteome is not saturated.

### AMPylated and Phosphorylated ATP Synthase Subunits

In further analysis, we focused on ATP synthase because it is a key enzyme which, apart from the synthesis of cellular energy - ATP - is directly involved in the mitochondrial mega-channel formation and the initiation of cell death [62–64]. Moreover ATP synthase dysfunctions due to mutations in mitochondrial and nuclear genes encoding its subunits lead to severe neurodegenerative, non-treatable diseases [65, 66]. The production of the most of ATP synthase subunits is induced upon respiration, the experimental conditions we applied (non-fermentative carbon source growth, [67]). The 27 AMPylation sites were found in seven enzyme subunits (Table S4 and Fig. S2), most in Atp2, Atp1 and Atp16. Among these AMPylation sites, absent in *fmp40Δ* mitochondria are: in Atp1 – T-366 and S-496, in Atp2 – T-124 and S-405, in Atp16 – S-29, T-42 and S-85, what makes them interesting modifications for further studies in the context of AMPylation by Fmp40.

Subsequently, we looked into the phosphorylation sites of ATP synthase subunits found in our experiments. In the literature, 102 phospho-sites were found while 129 were identified in enzyme subunits in our data (Table S4). 83 phosphorylation sites in ATP synthase subunits were not previously found (underlined in the last column of Table S4): 16 in Atp1, 27 in Atp2, 9 in Atp3, 9 in Atp4, 7 in Atp5, 5 in Atp7, 6 in Atp15, one in each Atp14, Atp18, Atp19 and Atp20. At the same time, the number of phosphorylation sites in ATP synthase subunits, that we did not detect in our data is 55 (underlined in the first column of Table S4). Thus, summarizing all data, there are as many as 140 phosphorylation sites in ATP synthase subunits. What is particularly interesting is that several positions in the three ATP synthase subunits are both AMPylated and phosphorylated: T-475 and Y-483 in Atp1, S-35, T-112, T-124, T-351, S-373, S-405 in Atp2, and S-29 in Atp16. This indicates a very sophisticated regulation of these subunits, especially Atp2, which appears to be the most extensively modified ATP synthase subunit.

### AMPylation of ATP synthase *in vivo*

We next aimed to determine the ATP synthase subunits AMPylation *in vivo* by Western blotting with the commercially available anti-Tre-AMP and anti Tyr-AMP antibodies. After the affinity chromatography of ATP synthase complex and subsequent separation of its subunits by SDS-PAGE, the proteins from the gel were transferred onto nitrocellulose membrane. Membrane was first incubated with anti-Tyr-AMP antibody and incubated with anti-ATP synthase subunits in order: Atp4, Atp5, Atp2, Atp16 and Atp3 (Fig. 3). Four quite strong bands and a fifth weaker one were detected by anti-Tyr-AMP. Two, in the region of 55 kDa correspond to Atp1 and Atp2 (see Fig. S1B to see the ATP synthase subunits separation in these experimental conditions stained by silver). Additional two strongly and one weakly decorated bands migrated et the region of 30 - 40 kDa. The Atp2 signal (after incubation with anti-Atp2 antibody) overlapped with one of ∼55 kDa bands, whereas Atp3 signal overlapped with the weakest band stained by anti-Tyr-AMP. On parallel the ATP synthase complexes were liberated from the inner membrane of mitochondria with the non-ionic detergent digitonin and separated in native gel. The lane fragments containing the monomers and dimers were subsequently put into the 2^nd^ dimensional denaturing 16 % gel and after migration proteins were transferred into the membrane. Membrane was first incubated with anti-Tyr-AMP antibody, then stripped, incubated with anti-Tre-AMP antibody, stripped again and incubated with anti-ATP synthase subunits in order: Atp4, Atp5, Atp2, Atp16 and Atp3 (Fig. S3). With this approach pattern of bands decorated by anti-Tyr and anti-Tre antibodies was same: two strong, migrating at ∼55 kDa, equivalent to Atp1 and Atp2, one strong migrating at 35 kDa, Atp3, and weakly decorated: one migrating above 15 kDa, non-identified protein, probably not ATP synthase subunit, one migrating below and only in the lanes from dimers – may be Atp20 or Atp21, and one migrating below which covers with Atp16. Incubation with anti-ATP synthase subunits identified and co-stained the anti-AMP signals of Atp1, Atp2, Atp3 and Atp16.

**Figure 3.**
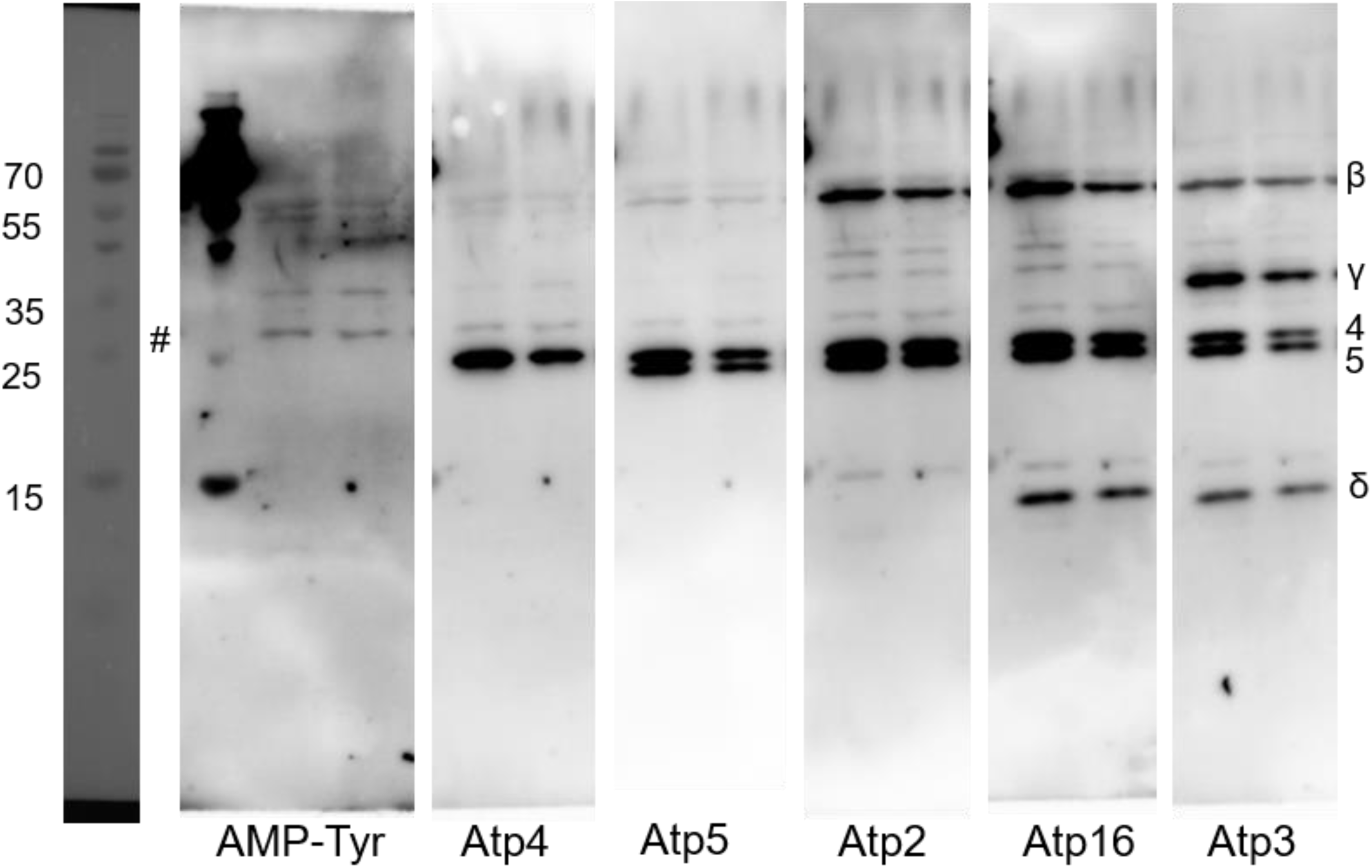
The AMPylation of ATP synthase subunits in the affinity purified enzyme trough Atp6-HA-6xHis. The whole ATP synthase complexes were purified from wild type strain mitochondria by Ni-NTA agarose beads and 2 µg of Ni-NTA bead eluates were loaded on the 15 % SDS-PAGE gel. After transfer, the membrane was incubated sequentially with the following antibodies: anti-AMP-Tyr, anti-Atp4,-Atp5,-Atp2,-Atp16 and-Atp3. Each lane contains an independent biological repeat. #- unknown protein, no ATP synthase subunit. The corresponding silver-stained gel is shown on Fig. S1B.

### Modification of the serine residue at position 29 in δ subunit of ATP synthase affects enzyme coupling

Within the mitochondrial ATP synthase, subunit δ (Atp16) plays a critical role in coupling the enzyme proton translocation and ATP synthesis activities. In yeast, deletion of the *ATP16* gene was shown to result in a loss of the mitochondrial genome resulting in 100% of petites cells (ρ^0^/ρ-, [68, 69]. This is a consequence of a passive proton transport through the enzyme proton channel, not coupled to ATP synthesis, and the loss of mitochondrial inner membrane potential. The loss of mtDNA is a rescuing event, preventing the synthesis of the two mtDNA-encoded subunits Atp6 and Atp9 forming the proton channel. We hypothesized that the regulation of the enzyme coupling at the level of subunit δ may be effectuated through the phosphorylation / AMPylation of serine residue at position 29, as this residue was found to be phosphorylated as well as AMPylated in our samples and is present at the end of the mitochondrial transit peptide of Atp16 as determined by AlphaFold structure prediction (Fig. 4A). To verify this assumption we introduced the codons for alanine or glutamic acid in the place of codon for serine 29 in the plasmid-borne *ATP16* gene. The plasmids were introduced into the wild type cells and *ATP16* gene sequence in the genome was subsequently replaced by the *KANMX4* gene – we assumed that the *ATP16–S29>A/E* variants will be functional and will not lead to the massive loss of mtDNA. In resulting strains the stability of mtDNA and the respiratory growth was checked. As shown in Fig. 4B, the Atp16–S29>A/E variants did not slowed down the growth of yeast cells on non-fermentable carbon source, in the presence of sub-inhibitory concentration of oligomycin as well, indicating the normal ATP synthesis rate. However the mtDNA stability in fermentative growth conditions was strongly decreased - in Atp16–S29>E cells culture the amount of petite cells reached 30% with 7% in the control cells and 12 % in Atp16–S29>A cells, suggesting the uncoupling of the proton translocation and ATP synthesis and the drop in the inner membrane potential (ΔΨ) (Fig. 4C). To check this we isolated mitochondria from cells grown in SC-glycerol-ura medium (due to the lack of growth disturbance on this carbon source) and measured the oxygen consumption and ATP synthesis rate. The oxygen consumption with NADH as an electron donor alone (state 4 respiration) was decreased of 10 and 20 % comparing to the control cells in Atp16–S29>A and Atp16–S29>E, respectively (Table S5). After successive addition of ADP (at the state 3, phosphorylating conditions) and CCCP (the uncoupled, maximal respiration) the similar decreases in oxygen consumption were seen (20 % and 30 %, respectively, Fig. 4D, Table S5). However, the rate of ATP synthesis by ATP synthase was similar as in the control cells. Thus, despite the full ATP synthase activity, the respiratory chain functions were reduced in the mutants. We conclude that S29 substitutions cause impaired signaling between ATP synthase and the respiratory chain, leading to increased ATP production per oxygen atoms consumed by respiratory chain (P/O ratios). We further investigated ATP synthase functionality in the mutant strains by membrane potential measurements, using Rhodamine 123, a cationic dye whose fluorescence is quenched when ΔΨ increases. For evaluating ΔΨ changes upon respiratory chain and ATP synthase activity, the mitochondrial membrane was first energized by feeding the respiratory chain with electrons from ethanol. The resulting ΔΨ was collapsed with complex IV inhibitor KCN and ATP was then rapidly added to. In these conditions the natural ATPase activity of ATP synthase inhibitor protein, IF1, is released from the enzyme and does not rebind [70]. The ATP addition induced in all strains mitochondria a large and stable ΔΨ that was fully reversed by oligomycin, thus dependent on ATPase activity of ATP synthase (Fig. 4E). Despite 100% of rho+ cells in these experimental conditions, we observed a slight (10%) decrease in the ΔΨ in both *atp16–S29>A/E* mutants, seen after addition of both the ethanol and the ATP. Based on the significant instability of mtDNA in *atp16–S29>A/E* mutants under growth conditions on a fermentable carbon source we hypothesized that the *atp16–S29>A/E* mutations cause a much higher perturbation of ΔΨ in cells grown on a fermentable carbon source. The isolation of mitochondria from yeast cells grown in glucose media is inefficient due to glucose repression of mitochondria biogenesis. We therefore measured the ΔΨ *in vivo* in cells grown on fermentative carbon source using the membrane potential-dependent fluorescent dye, MitoTracker Red, which, when used at the appropriate concentration, accumulates in mitochondria in an ΔΨ-dependent manner, such that the staining positively correlates with ΔΨ [50]. We used microscopic imaging to visualize and quantify ΔΨ. With microscopic imaging, using wild type cells treated with CCCP as a negative control, we were able to restrict the quantification of MitoTracker Red signal from mitochondria. As expected, we found that Atp16–S29>A/E cells exhibited lower ΔΨ of about 25 and 60% in Atp16–S29>A and Atp16–S29>E, respectively (Fig. 4F, Fig. S4). Therefore, the modification of the serine residue at position 29 of subunit δ, although it cannot be defined if AMPylation or phosphorylation, is directly related to the regulation of ATP synthase coupling under both fermentative and respiratory growth conditions (manifesting with decrease in ΔΨ) and the OXPHOS uncoupling under respiratory growth conditions (manifesting by the lack of increase of oxygen consumption at state 4 to compensate the drop in the ΔΨ).

**Figure 4.**
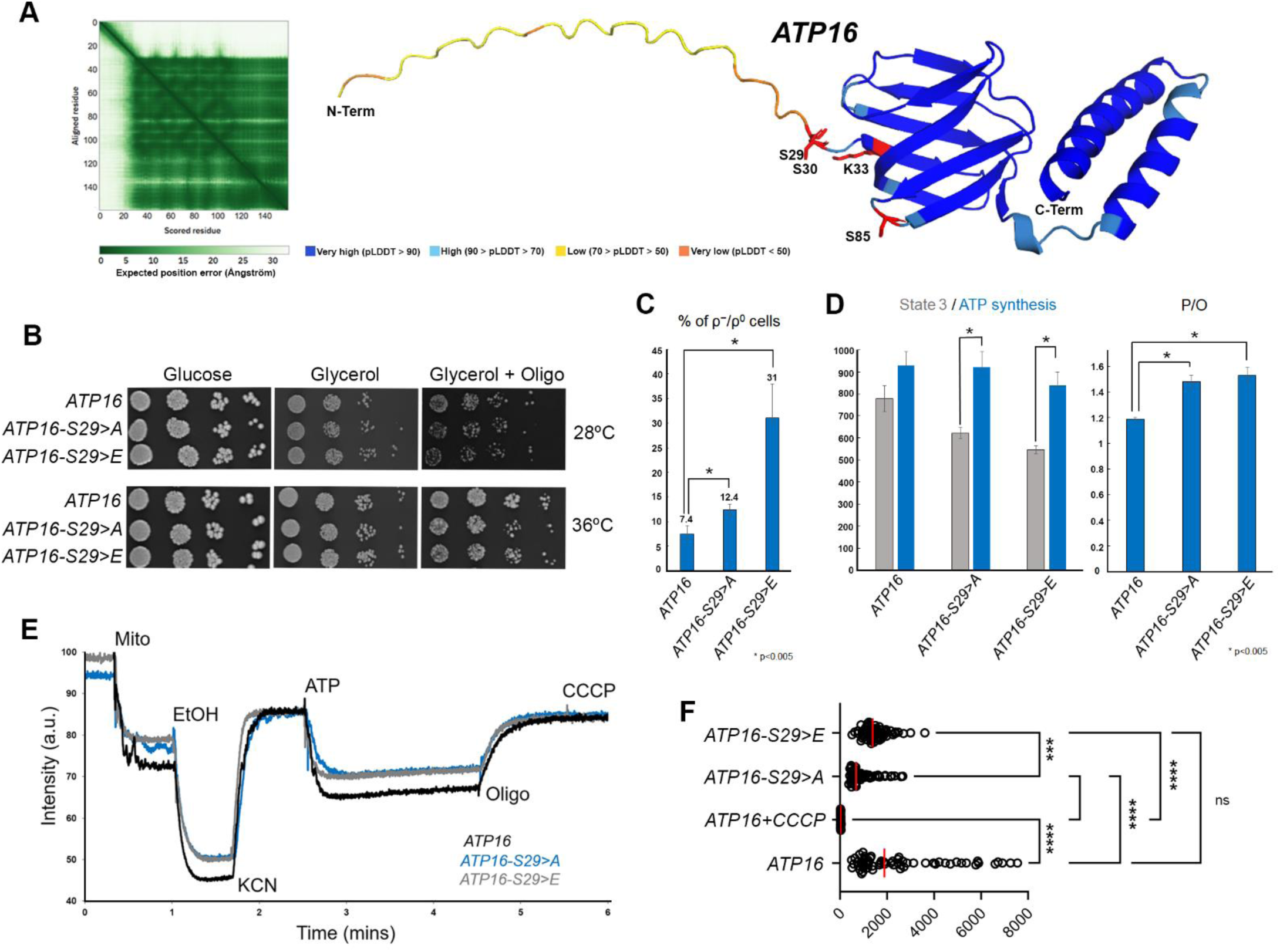
AMPylation / phosphorylation of serine residue in position 29 of subunit δ plays a role in ATP synthase coupling in both fermentative and respiratory growth conditions. **A** Position of S29 site in the AlphaFold predicted structure of Atp16 is marked in red. Other AMPylation sites in Atp16 identified in this study are indicated in red. **B** Changes in respiratory growth: cells from the indicated strains grown in SC-glucose-ura pre-cultures were serially diluted and spotted on rich glucose or glycerol plates with or without oligomycin (0.6 μg/mL) and incubated at 28 or 36°C. Plates without oligomycin were scanned after two days of incubation while those with the drug after seven days of incubation. **C** Stability of mtDNA: strains grown overnight in SC-glucose-ura were plated for single colonies on YPGA plates and replicated on rich glycerol plates to confirm the ρ^+^ vs ρ^−^/ρ^0^ genotype. The % of ρ^−^/ρ^0^ colonies were quantified. The graph presents data from analysis of two independent clones for each Atp16 variant, two biological and tree technical replicates for each strain. **D** Oxygen consumption at state 3 and ATP synthesis activities. The oxygen consumption was quantified in nmol O _2_·min^−1^ ·mg^−1^, the ATP synthesis in nmol of ATP.min^-1^.mg^-1^. **E** Variations in mitochondrial inner membrane potential in isolated mitochondria. The additions were 0.5 μg/ml of Rhodamine 123, 150 μg/ml of mitochondrial proteins (Mito), 10 μL ethanol (EtOH), 0.2 mM ATP, 2 mM potassium cyanide (KCN), 4 μg/mL oligomycin (oligo), and 4 μM carbonyl cyanide-m-chlorophenyl hydrazone (CCCP). **F** Fluorescence intensity in cells grown in glucose minimal selective medium and stained with of Mito Tracker CMXRos (see Materials and methods section for details). Statistical significance of the difference between the control *ATP16* expressing strain and ATP16 gene variants expressing strains and CCCP-treated negative control was determined using Kruskal-Wallis’ test in GraphPadPrism.The results are presented as a scatter plot of mean fluorescence intensity value for every individual count. Data from 60 cells per condition are pooled from four independent experiments. ***p =0.009 and ****p < 0.0001.

## Discussion

Although AMPylation is not a novel type of PTM in prokaryotes and eukaryotes our understanding of this modification is in its infancy, as both the number of identified AMPylases and the number of known substrate proteins is strikingly small [24, 34, 35]. In humans, the only AMPylases identified so far are HYPE and SelO proteins, and only the HYPE-dependent ampylation has been studied [25, 31, 36, 37]. The HYPE substrates list, identified in human chronic lymphocytic leukemia cell line by pulldown with antibodies against adenosine-phosphate and subsequently against AMP-Tyr, includes 213 proteins, while 25 proteins were screened in HEK293 cells in *in vitro* approach. In yeast one AMPylase was identified to date – the homologue of SelO, Fmp40 – and its substrates are not known, apart the Trx3, Prx1 and Grx2 mitochondrial redoxins [31, 34]. In this work we performed the first systematic profiling of AMPylation sites in proteins from isolated and percoll purified yeast mitochondria by label free quantitative MS approach, which does not distinguish low abundant AMPylated proteins from very abundant ounces. We identified 318 different AMPylation sites in 169 proteins. The most common and stable form of AMPylation occurs on the hydroxyl group of threonine, serine, or tyrosine through a phosphodiester bond and among 318 AMPylation sites, 174 modifications (55%) were present on those residues [71]. Less stable AMPylation modifications on lysine, histidine have been reported; however, as many as 100 modifications of lysine residues were detected in our data, which is quite significant (31 %) and indicates that it is a stable modification as well [72–74].

The identified AMPylated proteins cover a broad range of mitochondrial functions from bioenergetics: TCA cycle, oxidative phosphorylation, pyruvate and carbon metabolism, through redox proteins, mtDNA maintenance and translation, protein folding, transport to mitochondria, and amino acids and lipids biosynthetic processes, suggesting that many mitochondrial functions are regulated by reversible protein AMPylation. Apart from seven subunits of ATP synthase, four subunits of the ubiquinol-cytochrome c reductase complex, one subunit of cytochrome oxidase, eleven enzymes of TCA cycle were found to be AMPylated. Interestingly, none of the proteins involved in the communication of mitochondria with the nucleus were found. Proteins involved in cellular metabolism and translation were strongly represented among the HYPE substrates, beside proteins involved in maintenance of the cytoskeleton and cytoskeleton signaling, gene expression, ER-stress control, and redox homeostasis [36, 37]. A lot of mitochondrial enzymes were also found as a HYPE substrates in mammalian cells, while this enzyme is localized in the endoplasmic reticulum [75]. Among them are ATP synthase subunits α, β, g, e and δ, homologues of Pdx1 - PDHX pyruvate dehydrogenase protein X component or homologues of Nde1 and Ndi1 - NADH dehydrogenese subunits: NDUFA7, NDUFA12, NDUFB10 and NDUFS2, lactate dehydrogenase - homologues of which are on the list of yeast AMPylome. These results argue that the oxidative phosphorylation is regulated by AMPylation in mammalian cells as in yeast, however through different AMPylases.

We identified as many as 189 AMPylated peptides in cells lacking Fmp40. Although no homolog belonging to the FIC family in yeast is known [29], such a large number of Fmp40 independent modifications indicates on the presence of other AMPylases with specificity to serine, threonine and tyrosine, and to lysine and histidine. Modification of lysine residues in *E. coli* has been shown by AMP ligase, homologue of which in yeast is not identified [73, 74].

Studies of phospho-proteome of yeast cells have shown that the sub-fractionation of yeast extract to purify mitochondria significantly increased the sensitivity of detection of phospho-proteins [13, 17, 19, 21, 76–84]. The statistical analysis of the available data indicated that the detection of yeast phospho-proteins is approaching saturation [14, 84]. However we identified 1077 new phospho-sites in 516 proteins, including 823 new phospho-sites in 332 mitochondrial proteins. This number represents 2.4% of the phosphorylation sites already described [14, 17]. Taking into account that >50% of the phospho-sites were detected only by a single mitochondrial phospho-proteomic study like an observation that variation of phosphorylation sites in mitochondrial proteins is carbon sources influenced [19], we suggest that the yeast phospho-proteome can be expanded further.

Mitochondrial ATP synthase is subject to numerous PTMs. To date 102 phospho-sites were identified in 12 different enzyme subunits [13, 14, 19, 20, 85–88]. In our two approaches aiming to purify this enzyme we additionally identified 83 phosphorylation sites in ATP synthase subunits that were not previously found, which is as much as 45 % of all phospho-sites in ATP synthase. Significant amount (55) of previously identified ATP synthase phosphorylation sites were not detected in our study, confirming that each experiment is different and obtaining a complete phospho-proteome requires many approaches and repetitions. The reasons for this may be differences in the growth conditions, low stoichiometry of phosphorylation and other PTMs, and the LC-MS technical issues, as accessibility to tryptic digestion, poor phospho-peptide solubility or ionization efficiency and fragmentation due to neutral loss, due to which enrichment for phospho-peptides is advisable, which we did not apply here. The limits of informatics approaches for processing the mass spectrometry data also affect the level of detection [89–94]. Biological significance of Atp2 phosphorylation at T-124 or T-317 was demonstrated. The increased level of phosphorylated Atp2 resulted in higher abundance/activity of ATP synthase [20]. Mimicking phosphorylation at position 62 (S62>E) of Atp20 inhibited dimerization of the ATP synthase, whereas a block of phosphorylation by substitution of alanine for serine even enhanced the level of dimerization [13]. Our work extends the spectrum of ATP synthase modifications to include AMPylation. We report the presence of AMP moiety in 26 residues of seven ATP synthase subunits, while the Western-blot with anti AMP-Tyr or AMP-Tre antibodies decorated six bands. With high probability they are equivalent to Atp1, Atp2, Atp3, Atp16 and probably Atp15. The signal coming from the dimers only may be g/Atp20 or e/Atp21. Despite reproducing the experiment multiple times, we did not detect Atp20 or Atp21 AMPylation with MS further indicating that we did not detected all AMPylated proteins. This might suggest that modification rate is low, and more advanced MS methods (like FAIMS) or preparation (AMP enrichment) are necessary to achieve required sensitivity. The AMPylation sites were at the periphery of the subunits which is in agreement with a good accessibility to AMPylating enzyme/s (Fig. 5). Nine AMPylated positions were found phosphorylated as well (T-475 and Y- 483 in Atp1, S-35, T-112, T-124, T-351, S-373, and S-405 in Atp2, and S-29 in Atp16). AMPylation thus has potential to (reversibly or irreversibly) mask these phospho-sites, further substantiating a previously hypothesized crosstalk between these classes of PTM [95]. Among those doubly modified residues few were absent in *fmp40Δ* thus may be Fmp40-dependent (Atp1 – T-366 and S-496, in Atp2 – T-124 and S-405, in Atp16 – S-29, T-42 and S-85).

**Figure 5.**
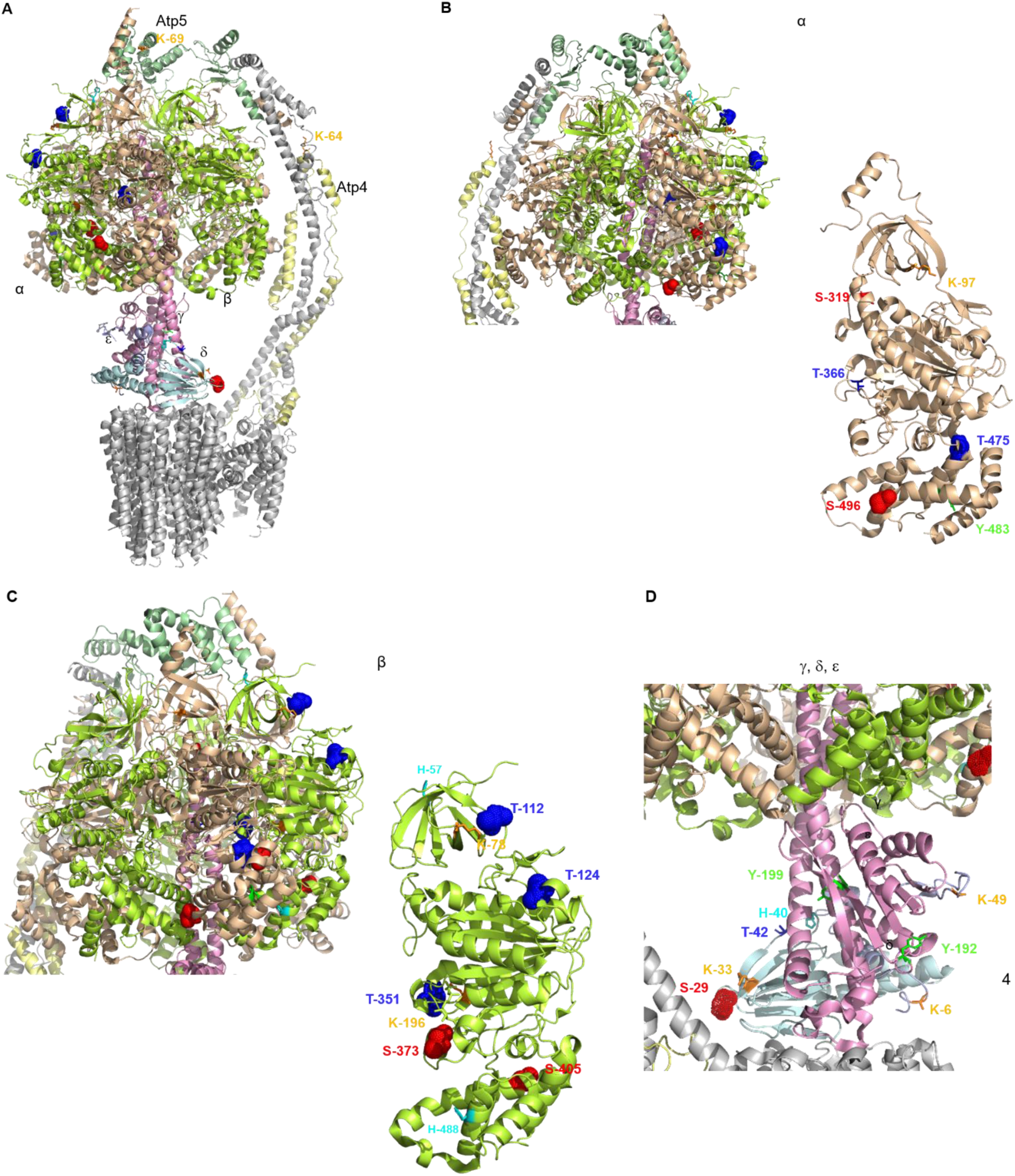
Distribution of AMPylated amino acid residues in the structure of ATP synthase subunits. **A** The overview of the ATP synthase monomer. **B** The F1 domain and modifications in subunit α. **C** The F1 domain and modifications in subunit β. **D** The central stalk subunits modifications. Subunits and modified residues are marked with colors, respectively, subunits: alpha – pale, beta – gray, Atp5 - pale green, Atp7 - pale yellow, gamma – light pink, delta – pale cyan, epsilon – light blue, residues: K – orange, T – blue, S - red, Y - green, H - cyan. AMPylated residues are shown as sticks while those found to be phosphorylated and AMPylated are indicated as spheres.

We demonstrated here the biological significance of Atp16-S29. Deletion of the *ATP16* gene was shown to result in a massive destabilization of the mitochondrial genome in the form of 100% of ρ^-^/ρ^0^ (petites) cells [68, 96]. Yeast cells lacking subunit δ cannot survive when are able to synthesize the ATP synthase proton channel due to lack of control of proton translocation by the ATP synthesis and the proton leak in consequence. Accordingly, the loss of mitochondrial DNA can be viewed as a rescuing event, as it prevents the synthesis of the two mtDNA-encoded subunits Atp6 and Atp9, forming the proton channel [69]. Thus the ATP synthase proton translocating activity is lethal to yeast cells missing the δ subunit. Our data show that the AMPylation or phosphorylation of serine residue in position 29 of subunit δ is directly related to the regulation of ATP synthase coupling (the control of proton flow through the channel in the membrane domain of the enzyme by the efficiency of ATP synthesis in the catalytic F1 domain of the enzyme) under both fermentative and the respiratory growth conditions. The loss of this control is compensated by ρ^-^/ρ^0^ mutations, observed in cells grown on fermentative carbon source. Although we did not observe an increase in the population of ρ^-^/ρ^0^ cells cultured on the non-fermentative carbon source, a 10% decrease in the inner membrane potential was found. This suggests that petite cells were eliminated from Atp16-S29>A/E cells cultures under respiration-dependent growth conditions. In contrast to cells lacking subunit δ, in which the decrease in the inner membrane potential is compensated by an increase in respiratory chain activity in state 4, in Atp16-S29>A/E cells this activity is reduced. Thus, the coupling of respiratory chain activity to ATP synthase activity, which in wild-type cells creates a potential on the inner mitochondrial membrane proportionally to ATP synthase activity [97], has been lost. Further experiments are needed to elucidate the mechanism of the OXPHOS coupling regulation by PTM modifications of ATP synthase subunits. This aspect and whether AMPylation of these residues depends on Fmp40 are currently under investigation in our laboratory.

## Conclusion

In contrast to bacterial AMPylation, which has gained renewed interest in recent years, eukaryotic AMPylation and its cellular substrates remain largely unexplored. Thus we are far from understanding the biological phenomena that may be regulated by this post-translational modification. The results presented here provide an initial view into the scope of AMPylation in yeast. Our findings open the door for extensive validation and research, establishing new directions in the field of eukaryotic AMPylation. Additionally, they provide a roadmap for substrate-targeted investigations that will elucidate the specific consequences of AMPylation on substrate structure, function, and involvement in disease.

## Data availability

The list of proteins identified in each mass spectrometry analysis from Database search using Mascot is available in **Supplementary Table S2 and S3**. The mass spectrometry proteomics data have been deposited to the ProteomeXchange Consortium via the PRIDE [98] partner repository with the dataset identifier PXD058585 and 10.6019/PXD058585“.

## Acknowledgements

We thank Prof. A. Tzagoloff for the MR6 ATP6-HisHA strain, Dr. Marie-France Giraud for the anti-ATP synthase subunits antibodies, Prof. Adrianna Skoneczna for providing deletion strains from Euroscarf Library. This work was supported by a grant from the National Science Center of Poland (2018/31/B/NZ3/01117) to RK. The permission number for work with genetically modified microorganisms (GMM I) for RK is 01.2-28/2017 – decision 72/2022. Fluorescence microscopy was performed in the Fluorescence Microscopy Facility of IBB PAS. The mass spectrometry experiments and analysis were performed in the Mass Spectrometry Facility of IBB PAS.

## Author Contributions

**CP:** Conceptualization; Data curation; Formal Analysis; Investigation; Methodology. **AW:** Data curation; Investigation. **SM:** investigation. **M.S.** Investigation. **KN:** Investigation. **EB:** Investigation. **DC:** Conceptualization; Data curation; Formal Analysis; Methodology. **AM:** Data curation; Formal Analysis. **RK:** Conceptualization; Data curation; Formal Analysis; Funding acquisition; Investigation; Project administration; Resources; Supervision; Validation; Writing – original draft; Writing – review & editing.

In addition to the <u>CRediT</u> author contributions listed above, the contributions in detail are: CP and RK conceived the project, CP designed all the experiments, performed the work, analyzed and interpreted the data; MS performed the growth tests. AW performed membrane potential measurements, KN and EB performed the pull-down of ATP synthase experiments, RK supervised the whole work, analyzed and interpreted the data, wrote the final manuscript. All authors worked on the manuscript.

---

**Table 1.**
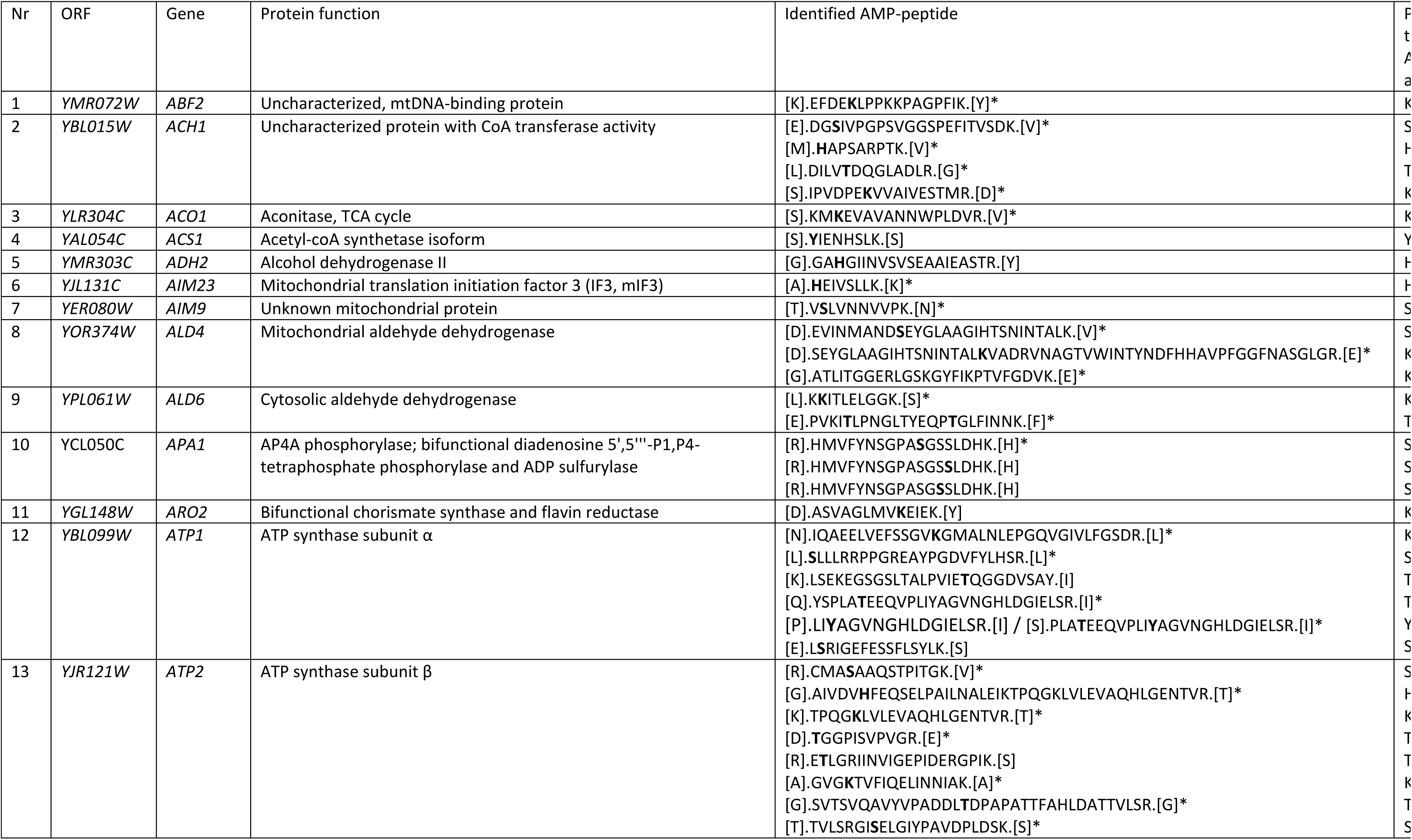
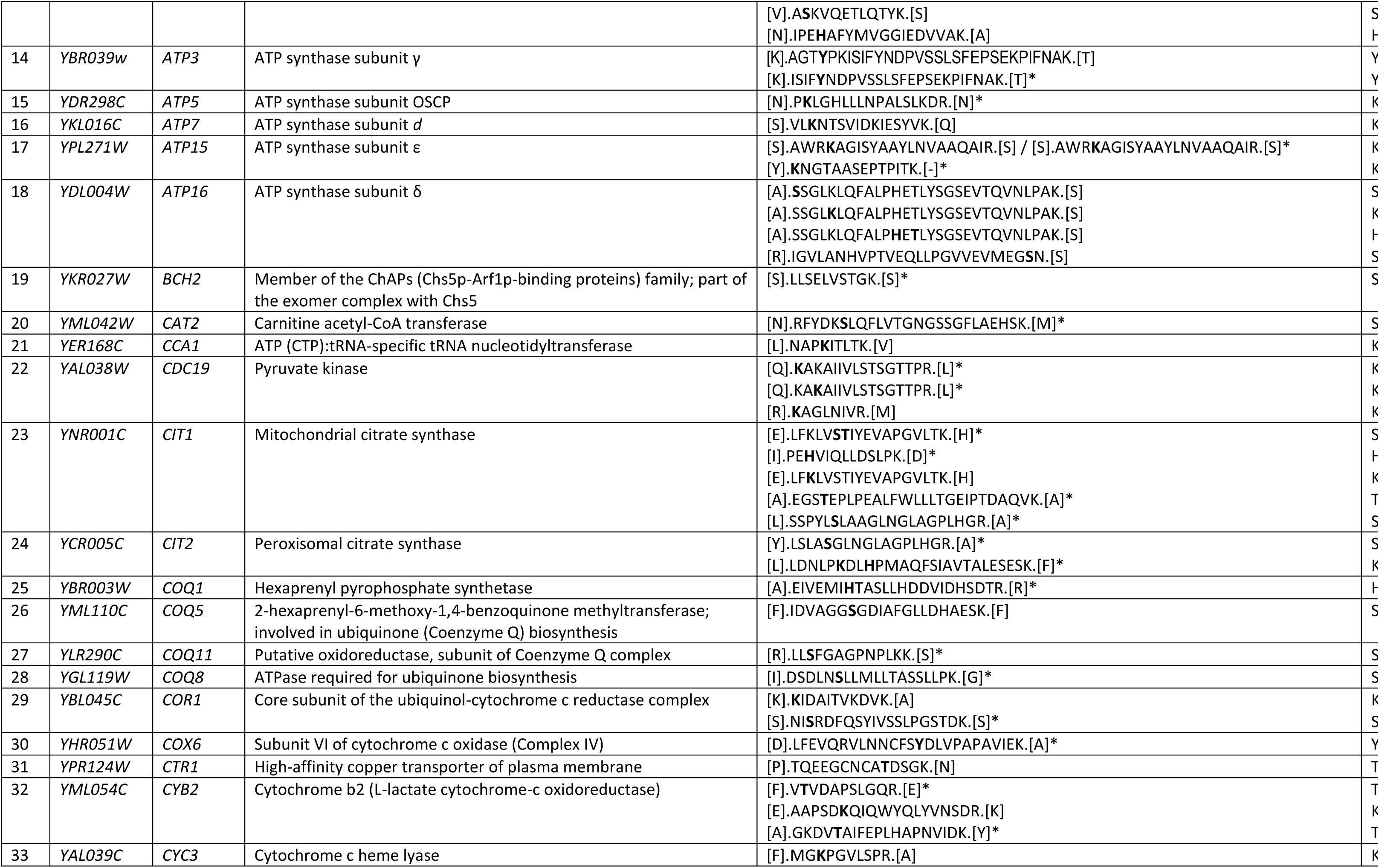
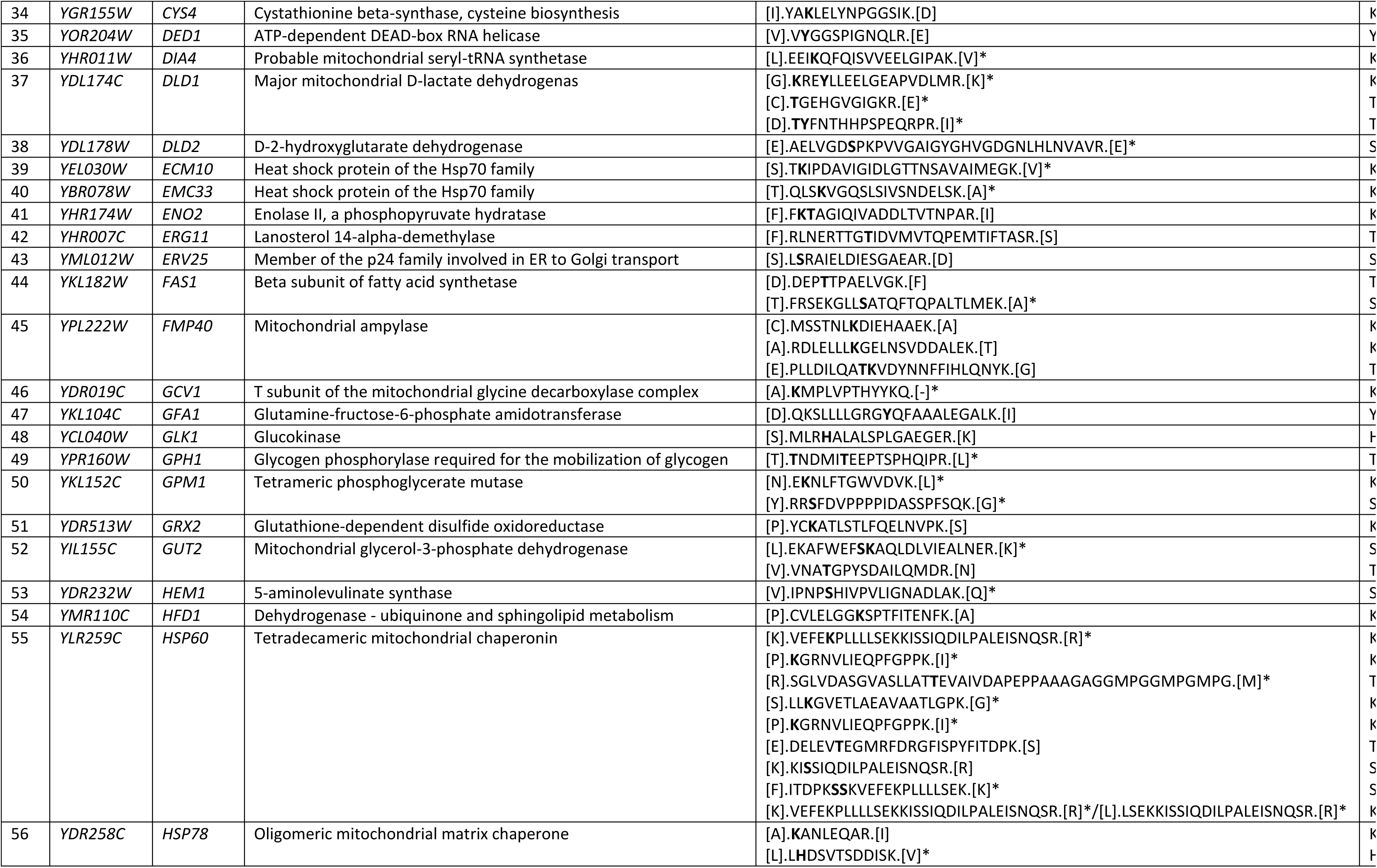
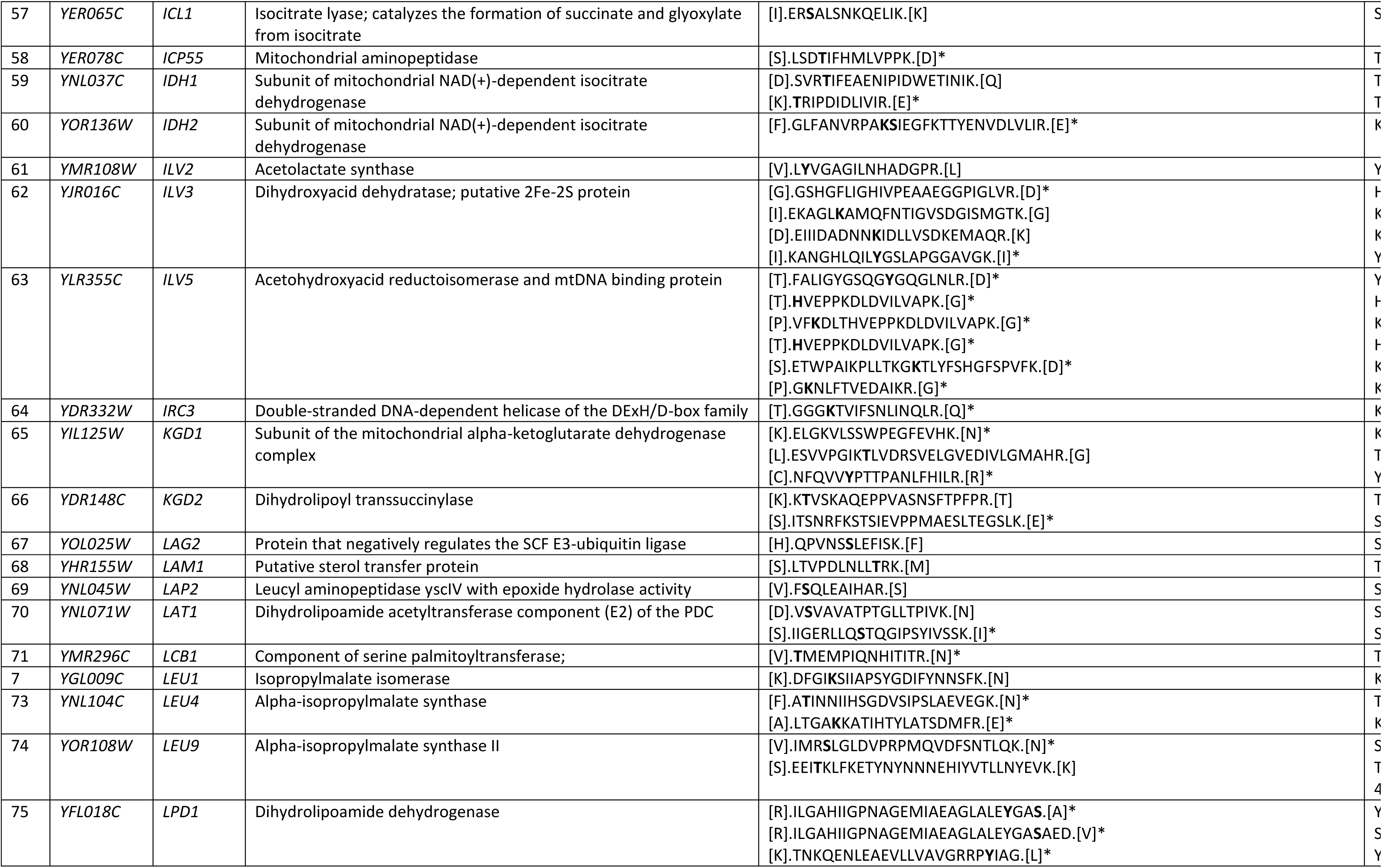
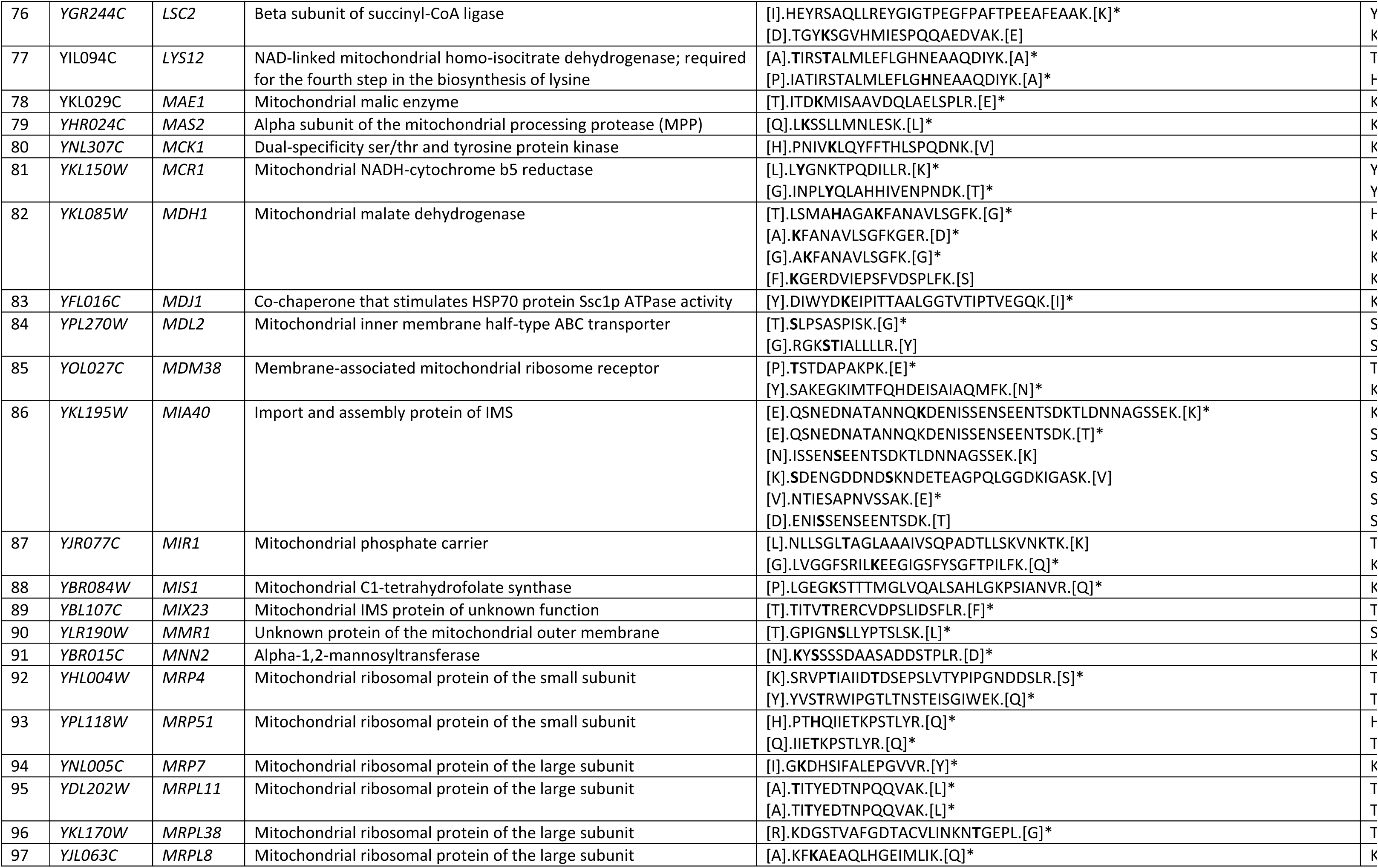
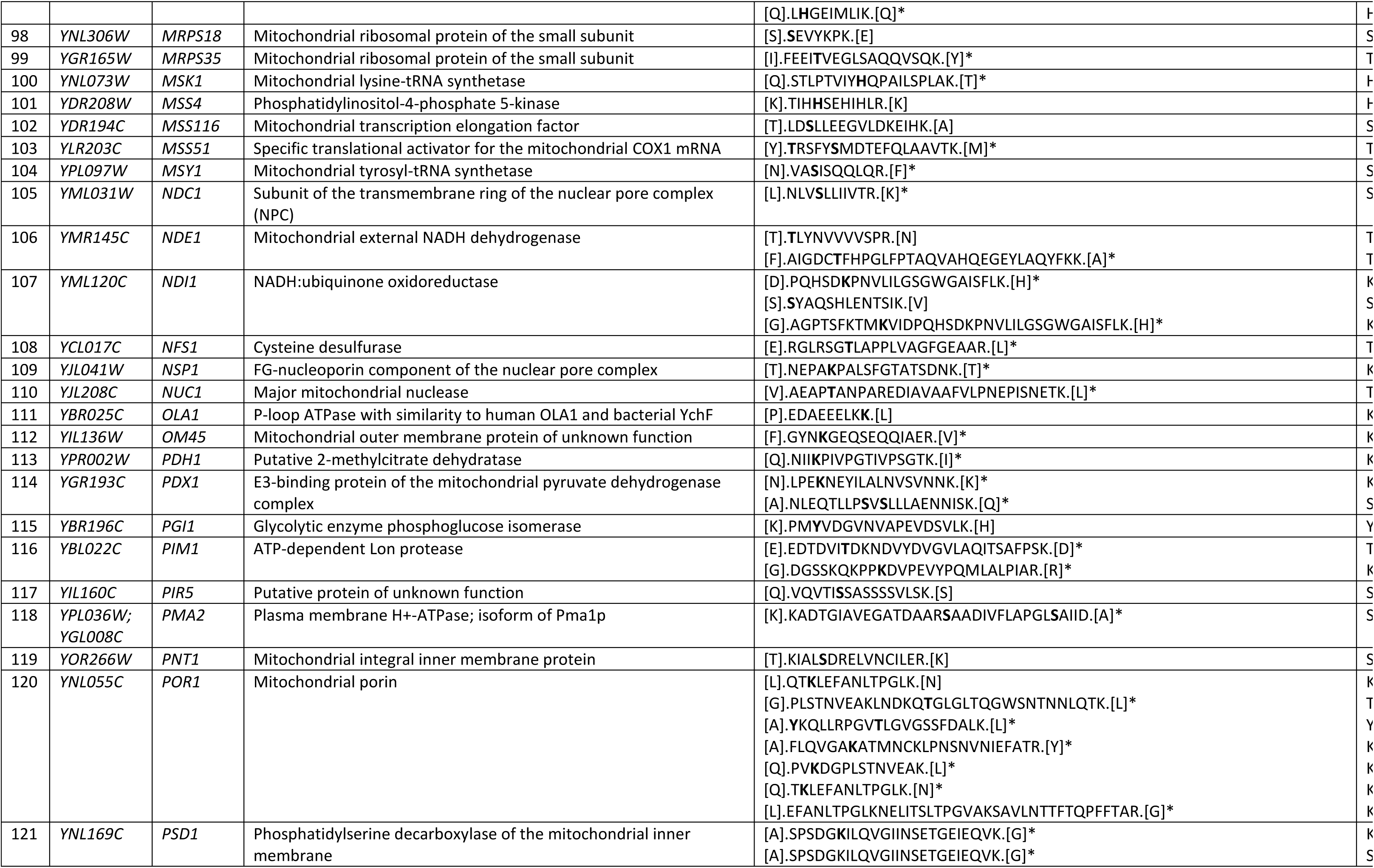
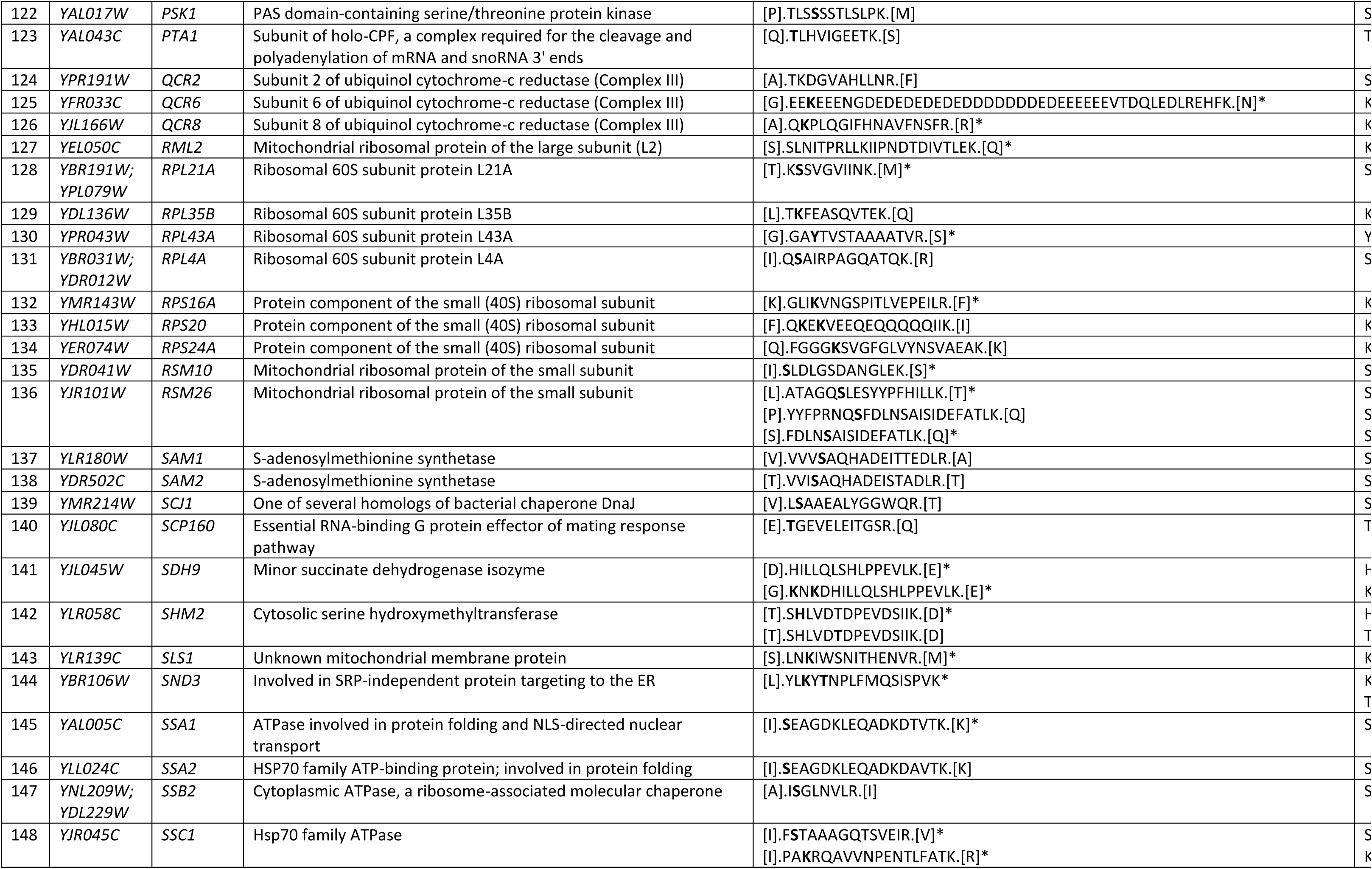
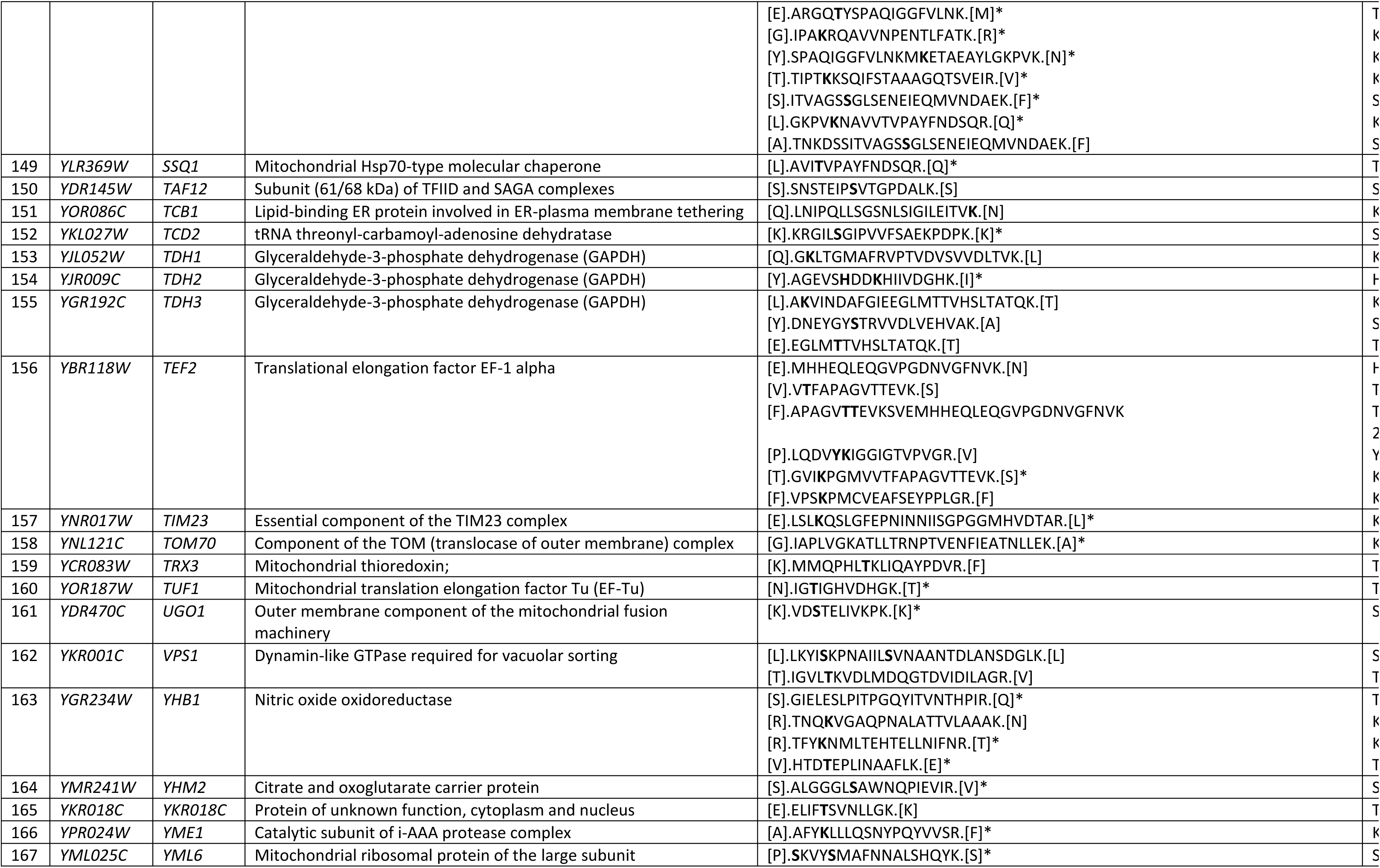
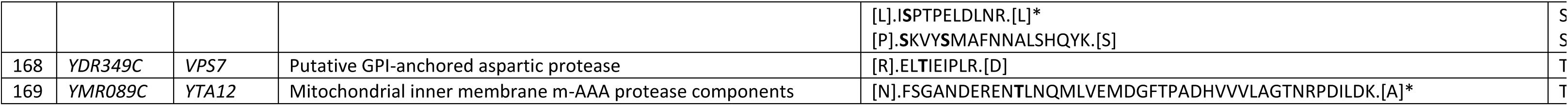
AMPylated peptides in purified yeast mitochondria identified in this study. *present in samples from *fmp40Δ*, AMPylated amino acid is indicated by bold font.

