## Supplementary Figures for "Profiling of yeast *Saccharomyces cerevisiae* mitochondrial AMPylome reveals a regulation of ATP synthase coupling trough subunit δ"

**Running title: *S. cerevisiae* mitochondrial AMPylome**

Chiranjit Panja, Suchismita Masanta, Aneta Wiesyk, Katarzyna Niedzwiecka, Emilia Baranowska, Dominik Cysewski, Marta Sipko, Anna Anielska-Mazur, Agata Malinowska, Roza Kucharczyk*
Institute of Biochemistry and Biophysics, Polish Academy of Sciences, Warsaw, Poland

### Supplementary Figures

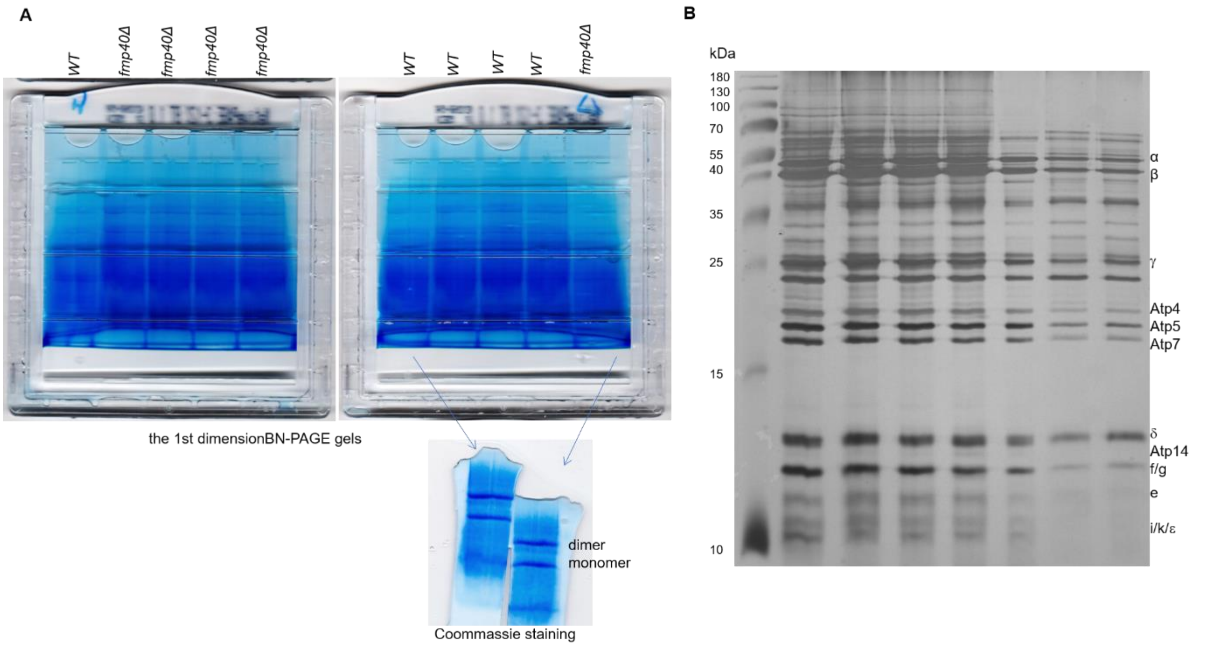


**Figure S1. A** The 1st dimension BN-PAGE gels separating the dimers and monomers of ATP synthase used for 2nd dimension SDS-PAGE or mass spectrometry analysis. The ATP synthase dimers and the monomers were extracted from 1 mg of the mitochondria with 1.5% of digitonin and run on NativePAGE™ 3–12 % Bis-Tris Gels. Gels were photographed before further processing. The indicated lanes were stained with Coomassie stain. **B** Silver stained gel separating the ATP synthase subunits after extraction of its complexes from the mitochondrial membrane and pull-down by Atp6-HA-6xHis. The eluates were separated in 15 % SDS-PAGE and the bands were visualized by silver staining. The dominating proteins identified in bands by mass spectrometry are indicated on the right. The representative silver stained gel is shown. Each lane contains an independent biological repeat.

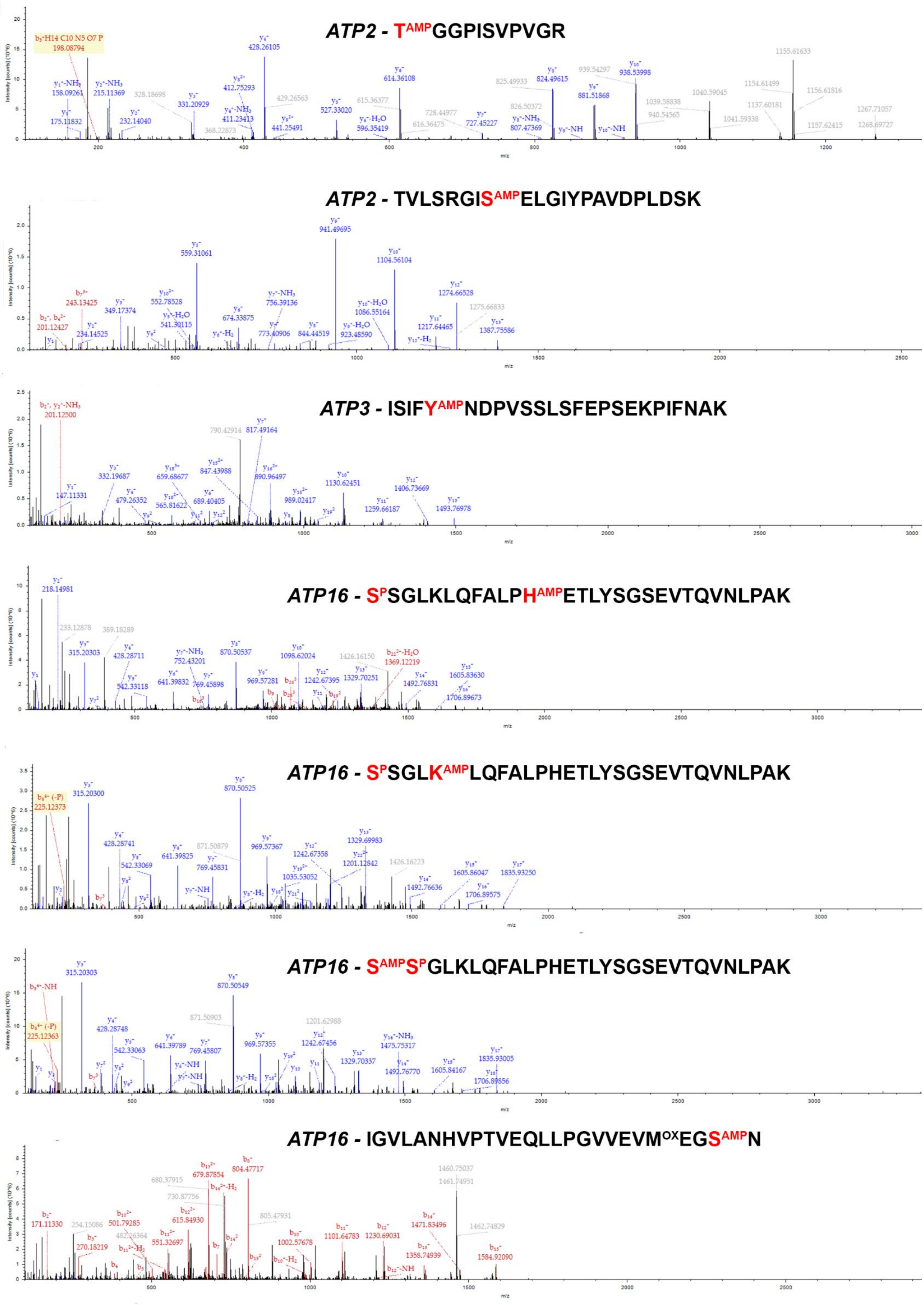


**Figure S2**. Spectra from Atp2, Atp3, and Atp16 are shown with the b-ions and y-ions used for identification marked. The post-translational modifications are highlighted in red in the representative peptides.

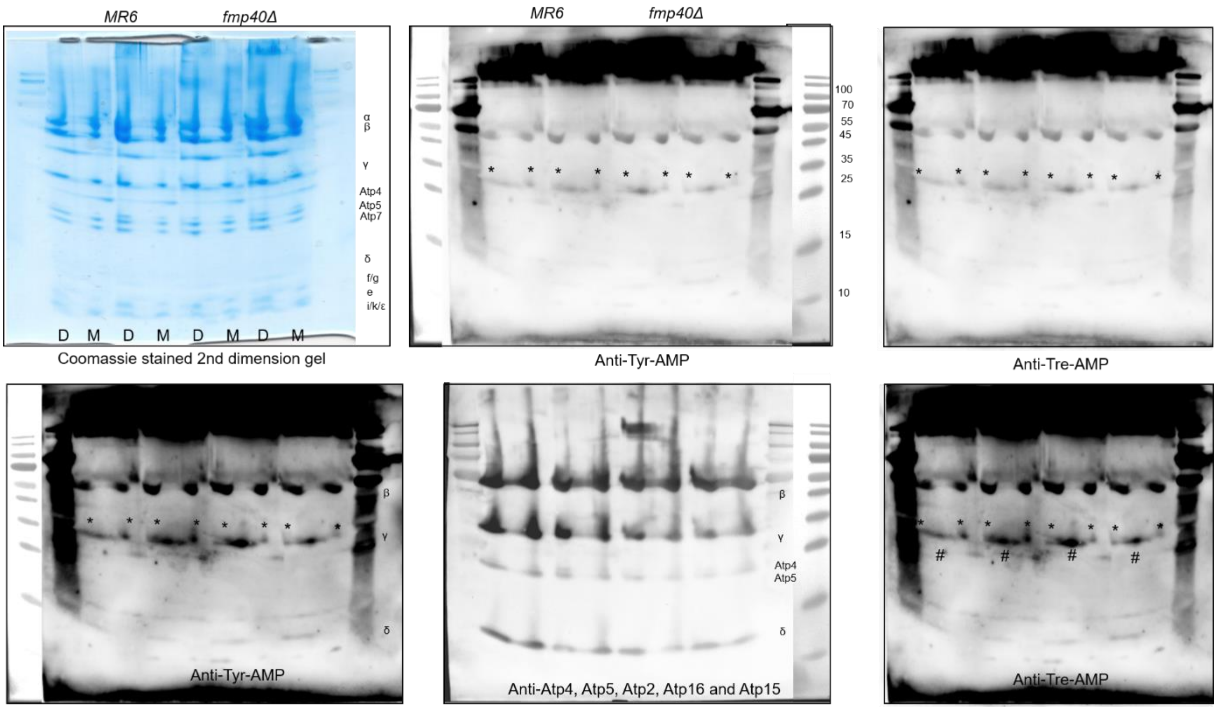


**Figure S3**. The AMPylated ATP synthase subunits *in vivo.* Second 2nd dimension gel after separation the dimers (D) and monomers (M) stained by Coomassie after the transfer of proteins to the membrane. Membrane was incubated with anti-Tyr-AMP, dehybridized, then with anti-Tre-AMP, dehybridized again and incubated subsequently with anti-ATP synthase subunits Atp2, Atp4, Atp5, Atp16 and Atp15. # - not identified protein from CIII or CIV, not ATP synthase subunit. * AMP signal seems to be absent/reduced in *fmp40Δ*.

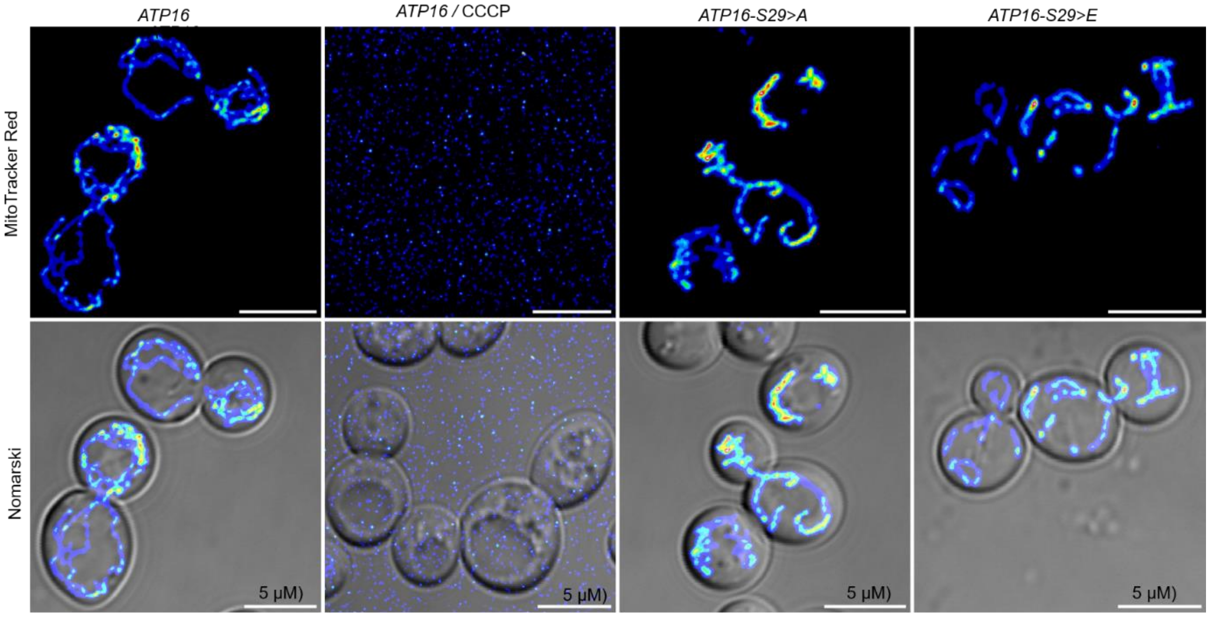


**Figure S4.** Representative images of wild-type, Atp16-S>A and Atp16-S>E cells stained with MitoTracker Red CMXRos. The CCCP-treated cells were introduced as a negative control. The images are representative of one of the four biological repetition from each strain. Scale bar represents 5 μm. Related to Fig. 4F.

**Table S5. Influence of the *atp16* mutations on yeast mitochondrial respiration and ATP synthesis.**

Mitochondria were isolated from wild type and mutants strains grown for 5-6 generations in SC-glycerol-ura at 28°C. Reaction mixes for assays contained 0.075 mg/ml of protein, 4 mM NADH, 150 (for respiration assays) or 750 (for ATP synthesis) µM ADP, 12.5 mM ascorbate (*Asc*), 1.4 mM N,N,N,N,-tetramethyl-p-phenylenediamine (*TMPD*), 4 µM CCCP. All strains contained 0% of $\rho^-/\rho^\circ$ cells. The values reported are averages of triplicate assays ± standard deviation. Respiratory and ATP synthesis activities were measured using freshly isolated, osmotically protected mitochondria buffered at pH 6.8. * p value < 0.05.

| Strain | Respiration rates, nmol $O.min^{-1}.mg^{-1}$ | | | | ATP synthesis rate, nmol $ATP/min^{-1}/mg^{-1}$ | P/O |
|---|---|---|---|---|---|---|
| | NADH | NADH +ADP | NADH +CCCP | Asc/TMPD + CCCP | | |
| WT | 283±24 | 779±59 | 1252±75 | 2381±263 | 928±63 | 1,19±0,01 |
| Atp16-S>A | 256±13 | 622±25 | 854±8 | 1703±75 | 920±71 | 1,48±0,05 |
| Atp16-S>E | 228±29 | 546±18 | 826±67 | 1663±94 | 838±62 | 1,53±0,06 |

*p<0,05
