## Supplementary TAble 4 for "Profiling of yeast *Saccharomyces cerevisiae* mitochondrial AMPylome reveals a regulation of ATP synthase coupling trough subunit δ"

Table S4. Modifications identified within ATP synthase subunits.

| Subunit (published P-sites)** | Identified AMP-peptide | Identified phospho-peptide | Position of the AMP / P amino a *** |
|---|---|---|---|
| Atp1/α | | [K].AQPTEVSSILEERIK.[G] / [K].AQPTEVSSILEER.[I] | T-43 |
| (S-37, | | [R].IKGVSDEANLNETGR.[V] / [K].GVSDEANLNETGR.[V] | S-57 |
| T-43, | | [V].FGLNNIQAEELVEFSSGVKGMALNLEPGQVGIVLFGSDR.[L] | S-93 |
| T-38, | | [V].FGLNNIQAEELVEFSSGVKGMALNLEPGQVGIVLFGSDR.[L] | S-94 |
| S-46, | [N].IQAEELVEFSSGVKGMALNLEPGQVGIVLFGSDR.[L] | | K-97 |
| S-47, | | [E].LVKRTGNIVDVPVGPGLLGR.[V] / [R].TGNIVDVPVGPGLLGR.[V] | T-128 |
| S-57, | | [R].RSVHEPVQTGLK.[A] / [R].RSVHEPVQTGLK.[A] / [R].RSVHEPVQTGLK.[A] | S-178 |
| T-128, | | [K].TAVALDTILNQKR.[W] / [K].TAVALDTILNQK.[R] | T-213 |
| S-178, | | [K].LYCVYVAVGQK.[R] | Y-237 |
| T-213, | | [K].LYCVYVAVGQK.[R] | Y-240 |
| S-230, | | [R].STVAQLVQTLEQHDAMK.[Y] / [R].STVAQLVQTLEQHDAMK.[Y] | S-248, T-249 |
| S-233, | | [R].STVAQLVQTLEQHDAMK.[Y] | T-256 |
| T-256, | | [F].TAASIGEWFR.[D] | T-286, T-289 |
| S-309, | | [K].HALIVYDDLSK.[Q] / [K].HALIVYDDLSK.[Q] | Y-305 |
| S-340, | [L].SLLLRRPPGREAYPGDVFYLHSR.[L] | | S-319 |
| S-350, | | [R].EAYPGDVFYLHSR.[L] / [L].SLLLRRPPGREAYPGDVFYLHSR.[L] | Y-331 |
| S-413, | [K].LSEKEGSGSLTALPVIETQGGDVSAY.[I] | | T-366 |
| S-445, | [Q].YSPLATEEQVPLIYAGVNGHLDGIELSR.[I] | [E].RLTQLLKQNQYSPLATEEQVPLIYAGVNGHLDGIELSR.[I] | T-462, T-475 |
| S-507, | | [Q].NQYSPLATEEQVPLIYAGVNGHLDGIELSR.[I] | Y-470, T-475 |
| S-511, | | [Q].YSPLATEEQVPLIYAGVNGHLDGIELSR.[I] | S-471 |
| S-527, | [P].LIYAGVNGHLDGIELSR.[I]/[S].PLATEEQVPLIYAGVNGHLDGIELSR.[I] | [Q].NQYSPLATEEQVPLIYAGVNGHLDGIELSR.[I] | Y-483 |
| S-533) | [E].LSRIGEFESSFLSYLK.[S] | | |
| | | | S-496 |
| | | [K].SNHNELLTEIR.[E] | S-511 |
| Atp2/β | [R].CMASAAQSTPITGK.[V] | [M].ASAAQSTPITGK.[V] | S-35 |
| (S-35, | [G].AIVDVHFEQSELPAILNALEIKTPQGKLVLEVAQHLGENTVR.[T] | | H-57 |
| S-39, | | [G].AIVDVHFEQSELPAILNALEIKTPQGKLVLEVAQHLGENTVR.[T] | S-61, T-74 |
| T-40, | [K].TPQGKLVLEVAQHLGENTVR.[T] | | K-78 |
| T-43, | | [R].TIAMDGTEGLVRGEK.[V] | T-94 |
| T-112, | | [T].IAMDGTEGLVRGEKVLDTGGPISVPVGR | T-100 |
| S-117, | [D].TGGPISVPVGR.[E] / [D].TGGPISVPVGR.[E] | VLDTGGPISVPVGR.[E] [K]. / [K].VLDTGGPISVPVGRETLGR.[I] | T-112 |
| T-124, | [R].ETLGRIINVIGEPIDERGPIK.[S] | | T-124 |
| T-237, | | [R].KPIHADPPSFAEQSTSAEILETGIK.[V] | S-156 |
| S-246, | | [A].DPPSFAEQSTSAEILETGIK.[V] / [K].LRKPIHADPPSFAEQSTSAEILETGIK.[V] | S-161,T-162,S-163 |
| S-299, | [A].GVGKTVFIQELINNIAK.[A] | | K-196 |
| S-302 | | [A].GVGKTVFIQELINNIAK.[A] / [L].FGGAGVGKTVFIQELINNIAK.[A] | T-197 |
| T-317, | | [K].AHGGFSVFTGVGER.[T] | S-215 |
| S-373, | | [R].EMKETGVINLEGESK.[V] / [K].ETGVINLEGESK.[V] | T-237 |
| T-470) | | [V].ALTGLTIAEYFR.[D] | T-268 |
| | | [R].FTQAGSEVSALLGR.[I] | T-295 |
| | | | S-310, Y-314, T-31[illegible] |

| | | | |
|---|---|---|---|
| | [G].SVTSVQAVYVPADDLTDPAPATTFAHLDATTVLSR.[G]<br>[T].TVLSRGISELGIYPAVDPLDSK.[S]<br>[V].ASKVQETLQTYK.[S]<br>[N].IPEHAFYMVGGIEDVVAK.[A] | [R].IPSAVGYQPTLATDMGLLQER.[I] / Q].PTLATDMGLLQER.[I]<br>[G].SVTSVQAVYVPADDLTDPAPATTFAHLDATTVLSR.[G] / [<br>[K].KGSVTSVQAVYVPADDLTDPAPATTFAHLDATTVLSR.[G]<br>[G].SVTSVQAVYVPADDLTDPAPATTFAHLDATTVLSR.[G]<br>[G].SVTSVQAVYVPADDLTDPAPATTFAHLDATTVLSR.[G]/[P].ADDLTDPAPATTFAHLDATTVLSR.[G]<br>[P].APATTFAHLDATTVLSR.[G] / [D].PAPATTFAHLDATTVLSR.[G]<br>[D].ATTVLSRGISELGIYPAVDPLDSK.[S]<br>[R].GISELGIYPAVDPLDSK.[S] / [I].SELGIYPAVDPLDSK.[S]<br>[P].LDSKSRLLDAAVVGQEHYDVASK.[V]<br>[R].LLDAAVVGQEHYDVASK.[V]/ [R].LLDAAVVGQEHYDVASK.[V]/<br>[A].VLEGKYDNIPEHAFYMVGGIEDVVAK.[A] / [N].IPEHAFYMVGGIEDVVAK.[A /<br>[A].VLEGKYDNIPEHAFYMVGGIEDVVAK.[A] [N].IPEHAFYMVGGIEDVVAK.[A] /<br>[K].YDNIPEHAFYMVGGIEDVVAK.[A] / [H].AFYMVGGIEDVVAK.[A] / | S-336,<br>T-338<br>Y-344<br>T-351<br>T-357, T-358<br>S-369, S-373,<br>S-373, Y-378<br>S-386, S-388<br>Y-401, S-405<br>Y-482, Y-491<br>H-488 |
| Atp3/γ (S-7, S-65, S-85, T-98, T-100, S-122, T-142, S-176, S-185, S-224, S-226, S-226) | [K].AGTYPKISIFYNDPVSSLSFEPSEKPIFNAK.[T]<br>[K].ISIFYNDPVSSLSFEPSEKPIFNAK.[T] | [K].ISAKKMDEAEQLFYK.[N]<br>[K].NAETKNLDVEATETGAPK.[E] / [K].NAETKNLDVEATETGAPK.[E]<br>[K].NLDVEATETGAPK.[E]<br>[I].VAITSDKGLCGSIHSQLAK.[A]<br>[K].GLCGSIHSQLAK.[A]<br>[R].HLNDQPNADIVTIGDK.[I] / [R].RHLNDQPNADIVTIGDK.[I]<br>[L].SINGIGKDAPTFQESALIADK.[L]<br>[K].DAPTFQESALIADK.[L]<br>[D].PVSSLSFEPSEKPIFNAK.[T]<br>[L].SFEPSEKPIFNAK.[T]<br>[K].TIEQSPSFGK.[F]<br>[K].TIEQSPSFGKFEIDTDANVPR.[D]<br>[K].FEIDTDANVPR.[D] | S-73<br>T-90<br>T-98<br>T-111<br>S-119<br>T-142<br>S-162<br>T-172<br>Y-192<br>Y-199<br>S-205<br>S-207<br>T-220<br>S-224, S-226<br>T-234 |
| *Atp4/b* (T-39, S124, S144, S-146, S-162, S-217) | | [Y].MSSTPEKQTDPK.[A] / [M].SSTPEKQTDPK.[A]<br>[K].ANSIINAIPGNNILTK.[T]<br>[K].AKANSIINAIPGNNILTKTGVLGTSA.[A]<br>[K].YLAPAYKDFADAR.[M]<br>[K].KVSDVLNASR.[N]<br>[R].IDSVSQLQNVAETTK.[V] / [K].DRIDSVSQLQNVAETTK.[V]<br>[K].DRIDSVSQLQNVAETTK.[V]<br>[K].ETVELESEAFELK.[Q]<br>A].KSVISRVQSELGNPK.[F] | S-38<br>S-52<br>T-64, T-66, T-71<br>Y-107<br>S-124<br>S-144, S-146<br>T-155<br>T-165<br>S-213 |
| Atp5 /OSCP (S-48, S-62, S-86, T-139, S-161, S-196, | [N].PKLGHLLLNPALSLKDR.[N] | [A].AAPPPVRLFGVEGTYATALYQAAAK.[N] / [A].PPPVRLFGVEGTYATALYQAAAK.[N]<br>[T].YATALYQAAAK.[N] / [T].YATALYQAAAK.[N]<br>[K].NSSIDAAFQSLQK.[V]<br>[K].LGHLLLNPALSLKDR.[N]<br>[K].DRNSVIDAIVETHK.[N] / [R].NSVIDAIVETHK.[N]<br>[C].FEKIASDFGVLNDAHNGLLK.[G] / [E].KIASDFGVLNDAHNGLLK.[G] / [K].IASDFGVLNDAHNGLLKG.[T] /<br>[E].KIASDFGVLNDAHNGLLK.[G] | T-35<br>Y-36, T-38<br>S-48, S-49<br>K-69<br>S-80<br>S-86, T-94<br>S-123 |

| | | | |
|---|---|---|---|
| T-199) | | [K].GTVTSAEPLDPK.[S]<br>[K].SLKLENVVKPEIK.[G] | T-139<br>S-169 |
| *Atp7/d*<br>(T-23,<br>S-25,<br>S-31,<br>S-32,<br>S-67,<br>S-95,<br>S-109,<br>S-112,<br>S-120,<br>T-121,<br>S-127) | [S].VLKNTSVIDKIESYV.[Q] | [K].VISSLRITGSTATQLSSFK.[K] / [R].ITGSTATQLSSFK.[K]<br>[R].ITGSTATQLSSFKK.[R]<br>[R].QLLELQSQPTEVDFSHYRSVLK.[N]<br><br>[S].VLKNTSVIDKIESYVK.[Q] / [K].NTSVIDKIESYVK.[Q]<br>[K].ETESLVSK.[E]<br>[K].DLQSTLDNIQSARPFDELTVDDLTK.[I] / [K].DLQSTLDNIQSAR.[P]<br>[K].DLQSTLDNIQSARPFDELTVDDLTK.[I]<br>[R].PFDELTVDDLTK.[I] | S-19, T-23<br>S-25<br>S-49<br>K-64<br>T-66, S-67<br>T-107<br>S-120, T-121<br>S-127<br>T-135 |
| Atp15/ε<br>(T-32,<br>S-34,<br>S-39,<br>T-41,<br>T46,<br>T-52) | [S].AWRKAGISYAAYLNVAAQAIR.[S]* /<br>[S].AWRKAGISYAAYLNVAAQAIR.[S]*<br><br><br><br><br>[Y].KNGTAASEPTPITK.[-] | <br><br>[S].AWRKAGISYAAYLNVAAQAIR.[S]<br>[R].SSLKTELQTASVLNR.[S]<br>[K].TELQTASVLNR.[S]<br>[R].SQTDAFYTQYK.[N]<br><br>[K].NGTAASEPTPITK.[-]<br>[I].VAITSDKGLCGSIHSQLAK.[A]<br>[L].SINGIGKDAPTFQESALIADK.[L] | K-6<br><br>S-10<br>S-24, S-25<br>T-28, T-32<br>S-39, T-41<br>K-49<br>T-52<br>T-111<br>S-162, T-172 |
| Atp16/δ<br>(S-21,<br>S-29,<br>S-30,<br>S-45,<br>S-47,<br>S-135,<br>S-136,<br>S-137) | [A].SSGLKLQFALPHETLYSGSEVTQVNLPAK.[S]<br>[A].SSGLKLQFALPHETLYSGSEVTQVNLPAK.[S]<br>[A].SSGLKLQFALPHETLYSGSEVTQVNLPAK.[S]<br>[A].SSGLKLQFALPHETLYSGSEVTQVNLPAK.[S]*<br>[R].IGVLANHVPTVEQLLPGVVEVMEGSN.[S] | [A].SSGLKLQFALPHETLYSGSEVTQVNLPAK.[S]<br>[A].SSGLKLQFALPHETLYSGSEVTQVNLPAK.[S] | S-29<br>S-30<br>K-33<br>H-40, T-42<br>S-85 |
| *Atp14/h*<br>(T-92,<br>T-120) | | [K].AYTEQNVETAHVAK.[E]* | T-86 |
| *Atp17/f*<br>S-17,<br>S-18,<br>S-23,<br>S-39) | | [K].SLPQGPAPAIK.[A] | S-39 |
| *Atp18/i*<br>*NONE* | | [K].AADLSSNTKEFINDPR.[N] | S-34 |

| | | | |
|---|---|---|---|
| *Atp19/kP-NONE)* | | [K].**T**VDIKTDNKDEEK.[F] | T-41 |
| *Atp20/g* (S-3, S-62) | | [K].Q**S**LNFALKP**T**EVLSCLK.[N] | S-62, T-70 |
| *Atp21/e* (S-90) | | [A].KLHPVV**T**PKDVPANASFNLEDPNIDFER.[V] | T-61 |
| *Mco10/l* (S-45, S-46, S-49, S-53, S-56, S-57, S-59, S-71) | | [K].**S**GE**S**SNSDAMGKDDDVVK.[S]<br>[K].**S**IEGFLNDLEKDTR.[Q] | S-53,S-56<br>S-71 |

*a* Phosphorylated / ampylated amino acid is indicated by blue / red color, respectively. Both AMPylated and phosphorylated residues are marked in both colors.

*b* Positions were determined from precursor protein sequences.

*- present in *fmp40Δ*

** - X – not found in our conditions

*** X – new phosphorylation site found
